# FMO and CYP monooxygenase families determine the metabolic flux of hydroxylated tryptamine derivatives in barley (*Hordum vulgare*) following pathogen infection

**DOI:** 10.1101/2025.05.25.656007

**Authors:** Joachim Møller Christensen, Mette Marie Toldam-Andersen, Hans Jørgen Lyngs Jørgensen, Nikola Micic, Nanna Bjarnholt, Mohammed Saddik Motawie, Mallika Vijayanathan, Mathilde Pizaine, Jan Günther, Louise Kjærulff, Dan Stærk, Meike Burow, Sara Thodberg, Elizabeth H J Neilson

## Abstract

To counteract pathogenic microorganisms, plants execute a complex resistance response that includes major metabolic reprogramming and production of bioactive defensive compounds. Barley (*Hordeum vulgare*) is a major cereal crop, but suffers significant yield losses due to pathogen attack every year. Here we use an untargeted metabolomic approach to assess the diversity and shifts in key barley metabolites produced in response to *Pyrenophora teres f. teres* infection, a hemibiotrophic fungal pathogen and causal agent of net blotch disease. Tryptophan-derived compounds, including tryptamine, serotonin, and a novel indole alkaloid – 2-oxo-tryptamine (2OT) – were among the most significantly induced and abundant compounds, with mass spectrometry imaging revealing that these metabolites accumulate at the site of infection. A transcriptomic approach identified a flavin-containing monooxygenase (FMO), which was functionally characterized as a 2-oxo-tryptamine synthase (2OTS). In addition, a cytochrome P450 (CYP71P10) was characterized as a tryptamine-5-hydroxylase, responsible for serotonin biosynthesis. These characterized genes are tightly co-expressed with genes involved in tryptophan biosynthesis and signify a major metabolic flux towards indolic compounds after infection, potentially serving as bioactive phytoalexins. Additional microbial interactions using a biotrophic fungus (*Blumeria hordei*; powdery mildew), a hemibiotrophic bacterium (*Pseudomonas syringae*), and alternative cultivars suggest that these pathways represent a generally activated resistance response in barley. These results provide new insights within the barley defense response relevant for the development of disease resistant traits in cereal crops.

## Introduction

Barley (*Hordeum vulgare* L.) is the fourth most cultivated cereal crop, for food, fodder and raw materials. An estimated average 15% of barley yield is lost per annum due to pathogens (Oerke & Dehne, 2004), but this loss can be up to 40% depending on the pathogen and environmental condition (Clare et al., 2020). Global warming alters the range of phytopathogens to which plants are exposed, as well as their susceptibility towards these pathogens. Consequently, climate change leads to an overall increase in disease pressure (Sangiorgio et al., 2020; Seth & Sebastian, 2024). Coupled with the drastically increasing world population, estimated to reach 9 billion by 2050, and the associated rise in food demand, there is an urgent need to expand our knowledge on plant-pathogen interactions at both molecular and systemic levels (Raza et al., 2019).

Plants produce a large array of bioactive specialized defense metabolites that provide chemical protection against pathogen attack. Plant defense metabolites are broadly categorized into phytoalexins and phytoanticipins (VanEtten et al., 1994). Phytoalexins are synthesized *de novo* in response to biotic challenges such as pathogen attack or herbivore infestation, while phytoanticipins are constitutively present in plant tissues, acting as preemptive protective agents. Barley produces many different specialized metabolite classes, including benzofurans, hydroxynitrile glucosides (HNGs), terpenoids, phenylamides, and alkaloids; each with varying modes of action. For example, benzofurans (i.e. hordatines) possess antifungal activity and accumulate in the seedling leaves (Hamany Djande et al., 2022), providing early-stage protection (A. Stoessl & C. H. Unwin, 1970), but can also be inducible in mature plants (Laupheimer et al., 2023). Similarly, levels of barley-specific diterpenes (i.e. hordedanes) are induced after infection and exuded from barley roots, exhibiting both antifungal activity, and modify fungal root colonization (Liu et al., 2024). Tryptophan-derived metabolites represent a further metabolite class, highly relevant for defense in barley and other important cereal crops. Specifically, the tryptophan-derived metabolites tryptamine, serotonin and phenylamides (e.g. triticamides) consistently accumulate in barley after infection by a wide range of pathogenic hosts (Lemcke et al., 2021; Ube, Harada, et al., 2019; Ube, Yabuta, et al., 2019). Barley also produces the tryptophan-derived indole-alkaloid gramine, 5,5′-dihydroxy-2,4′-bitryptamine (DHBT; a dehydrodimer of serotonin), and 3-(2-Aminoethyl)-3-hydroxyindolin-2-one (AEHI; di-hydroxylated tryptamine derivative); for all of which, bioactivity has been reported (Corcuera, 1984; Ishihara et al., 2017; Lu et al., 2018; Wippich & Wink, 1985).

Despite significant progress towards identifying and measuring the chemical defense constituents produced by barley in response to pest and pathogen attack, biosynthetic pathway elucidation within barley is limited to a few compound classes: hydroxynitrile glucosides (Knoch et al., 2016), hordedane diterpenes (Y. Liu et al., 2024) and the indole alkaloid gramine (Dias et al., 2024; Ishikawa et al., 2024; Leland & Hanson, 1985). Transcriptomic studies show significant upregulation of genes leading to tryptamine biosynthesis, but downstream assessment of where this metabolic flux leads, is currently unknown (Lemcke et al., 2021). In contrast to other members of the Poaceae family, cultivated barley (i.e. *H. vulgare*) does not produce the indole-derived defense class of benzoxazinoids (BXs) (Ube et al., 2017). In other grasses, the core BX pathway initiates by the formation of indole by the action of an indole-3-glycerol phosphate lyase (BX1), a diversified homologue of the alpha subunit of tryptophan synthase (Zhuang et al., 2012). Indole then undergoes four consecutive hydroxylations leading to the first considered benzoxazinoid (i.e. DIBOA) by four different CYP71C enzymes (BX2-5), co-occurring in a tightly regulated gene cluster (Frey et al., 1997; Wu et al., 2022). In a case of convergent evolution, the first two hydroxylations (i.e. BX2 and BX3) are catalyzed by Class B flavin-containing monooxygenases (FMO) in the BX-producing eudicots *Consolida orientalis*, *Lamium galeobdolon* and *Aphelandra squarrosa* (Florean et al., 2023, 2025). While lacking BXs, barley still possess orthologous *BX2-4* genes (Sue et al., 2011), as well as several related FMOs, that are significantly induced after infection (Sjokvist et al., 2019). In the absence of a *BX1* gene (and BX’s) it is speculated that barley has evolved a defense pathway that is channeled towards tryptophan-derived metabolic defense response in contrast to indole-derived metabolites observed for other cereals.

In this study, we subject the elite malting barley cultivar RGT Planet to infection by *Pyrenophora teres f. teres* (henceforth *P. teres*) infection (the causal agent of net blotch), to characterize its metabolic response. *Pyrenophora teres* is a major hemibiotrophic fungal pathogen that can result in up to 40% yield loss in barley (Clare et al., 2020). Untargeted metabolomic analysis identified tryptamine, serotonin and two novel hydroxylated tryptamine derivatives to be significantly induced after *P. teres* infection, with the most abundant hydroxylated tryptamine isomer identified as 2-oxo-tryptamine (2OT). A combination of metabolomics, transcriptomics and heterologous expression in *Nicotiana benthamiana* and *Escherichia coli* identified an FMO from the N-OX3 family as an enzyme that catalyzes the production of 2OT from tryptamine, and hereby name it 2-oxo-tryptamine synthase (2OTS). Furthermore, we biochemically characterize CYP71P10, a tryptamine 5-hydroxylase, as the serotonin-producing gene in barley. These results suggest that the rapid induction of tryptamine-derived metabolites constitutes an important metabolic branch of the barley defense response to pathogen attack.

## Results

### Disease progression of net blotch in barley

In this study, we investigated metabolomic changes induced in barley *cv.* RGT Planet after infection with the hemibiotrophic fungal pathogen *P. teres*, the causal agent of net blotch – a major foliar disease of barley responsible for tremendous yield loss and decrease in seed quality worldwide (Tini et al., 2022).

Progression of disease symptoms were observed visually for non-treated, control and infected leaves at 0, 1, 2, 4 and 7 days post inoculation (dpi). At these timepoints, samples were also collected in five biological replicates for metabolomic and transcriptomic analyses. Clear signs of infection were noticeable in the pathosystem (Figure 1). Around 2 dpi, phenotypical signs of fungal infection were already visible with infected leaves appearing more yellow compared to control and by indications of the longitudinal necrotic lesions, which are characteristic for net blotch (Tini et al., 2022). At 4 dpi, infected leaves were clearly affected with brown necrotic lesions surrounded by chlorotic areas. Hereafter, net blotch symptoms developed rapidly with chlorotic areas increasing in size leading to wilting and leaf death at 7 dpi. No disease symptoms were detected on the control leaves (treated with water), which remained healthy and green throughout the experimental time course. With the aim of investigating the metabolic reconfiguration in the susceptible barley cultivar throughout disease development, an untargeted metabolomic approach was applied.

**Figure 1:**
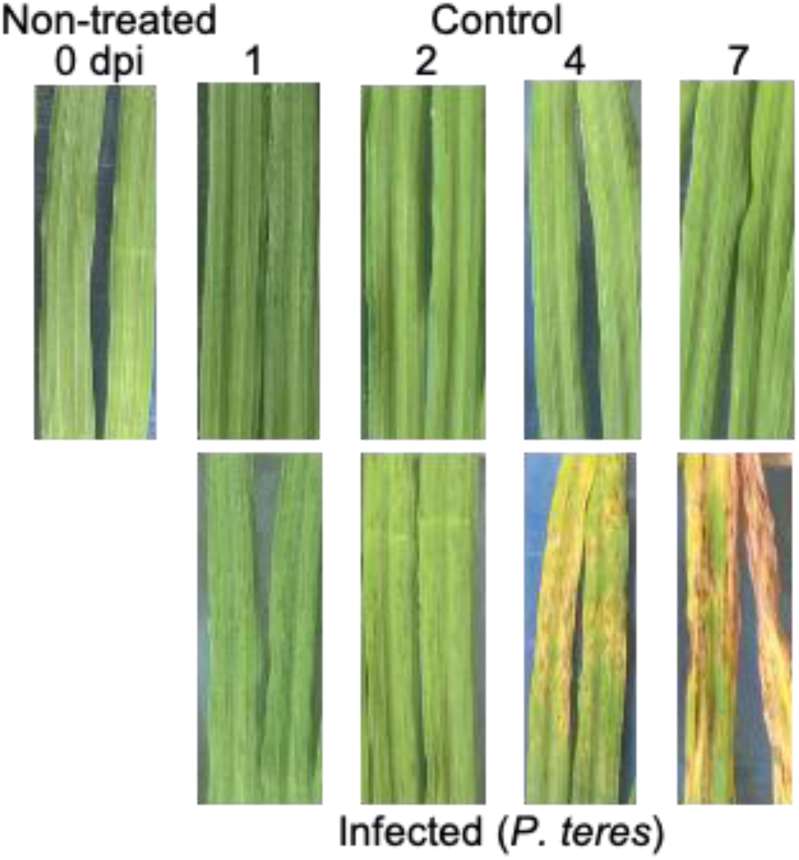
Disease progression of net blotch on barley *cv.* RGT Planet leaves inoculated with *P. teres*. Development of disease symptoms at 0 (non-treated), 1, 2, 4 and 7 days post inoculation (dpi). Representative pictures of infected and control leaves.

### Chemical Profiling of Barley Leaves Following Infection by the Fungus *Pyrenophora teres f. teres*

An untargeted metabolomics approach was used to investigate the chemical profile of barley leaves in response to *P. teres* infection. Leaf methanol extracts were analyzed by UHPLC-qTOF-MS/MS and subsequently processed in MZmine4. The generated feature table and spectral files were used as input for various metabolomic computational tools, including feature-based molecular networking (FBMN), SIRIUS and CANOPUS (Figure 2A). Prior to MZmine4 data processing, overall reliability of the spectral data acquisition throughout the analysis was assessed and confirmed by principal component analysis (PCA), which showed narrow clustering of quality control (QC) pooled samples and blanks (Figure S1).

**Figure 2:**
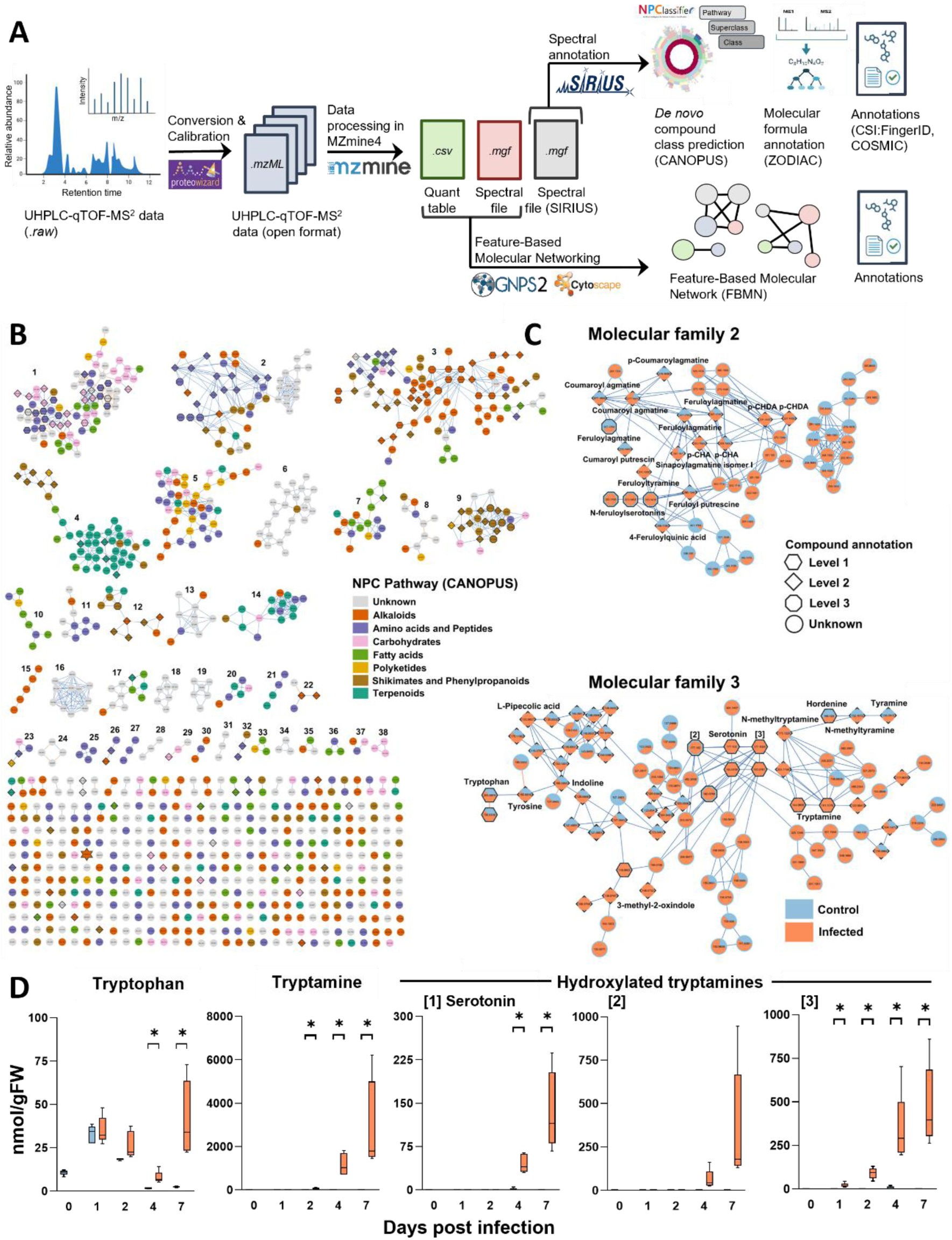
Chemical profiling of barley leaves after fungal infection by *P. teres.* **(A)** Data processing workflow used for the metabolite profiling and annotation of the acquired spectral data set from barley leaf methanol extracts. **(B)** Global feature-based molecular network (FBMN) visualizing the detected chemical space of barley *cv.* RGT Planet. The 38 largest molecular families (i.e. ≥ 3 nodes) in the network are numbered. Nodes are colored according to chemical NPC pathway predictions by CANOPUS. The shape corresponds to the level of compound annotation. The star represents AEHI (level 1). **(C)** Enlarged depiction of molecular families 2 and 3 highlighting features (primarily phenylpropanoids and alkaloids) that are highly induced following *P. teres* infection. Pie-charts indicate relative abundance of individual features in control (blue) and infected (orange) samples across entire spectral dataset. **(D)** Abundance of tryptophan, tryptamine, and three hydroxylated tryptamine derivatives, which are strongly induced after pathogen inoculation. Abbreviations: AEHI, 3-(2-aminoethyl)-3-hydroxy-indolin-2-one (Ishihara et al., 2017); NPC, NPClassifier taxonomy (Kim et al., 2021); *p*-CHA, *p*-coumaroyl-hydroxyagmatine; *p-*CHDA, *p*-coumaroyl-hydroxydehydroagmatine.

FBMN compares and distinguishes between chromatographic features, which are then connected to one another based on spectral similarity. Each detected feature is depicted as a node in the network, and related nodes are grouped in molecular families (MFs) (Nothias et al., 2020). The generated FBMN of the entire barley spectral dataset (non-treated, control, and infected leaves) comprised 998 nodes, each representing detected metabolite features with MS2 spectral information. In the network, 631 nodes clustered into 75 MFs (2 or more connected nodes) based on their spectral similarity (Figure 2B). The remaining 367 nodes were represented as singletons, i.e. not included in a MF.

76 features were removed from the generated feature table during blank subtraction resulting in a total of 922 features. Semi-automated putative annotation of 106 metabolites was achieved with confidence level 2 (Schymanski et al., 2014; Sumner et al., 2007) by matching (> 0.7 cosine score) the processed LC-MS/MS data output to the publicly available spectral libraries, GNPS and MoNA. This annotated metabolite list corresponds to an annotation rate of approximately 12% of the detected features. The metabolite annotation rate was further improved to 14% by matching 25 features to in-house available authentic standards (confidence level 1) analyzed with the same LC-MS/MS method as the samples. Additionally, annotated features in the molecular network could in some cases be used as anchoring nodes to propagate the information and manually annotate structurally similar metabolites in the same MF by comparison to spectral data reported in literature (confidence level 2), e.g., *p*-coumaroyl-hydroxyagmatine (*p*-CHA) and *p*-coumaroyl-hydroxydehydroagmatine (*p*-CHDA) in MF 2 (von Röpenack et al., 1998) and a methoxychalcone in MF 12 (Ube et al., 2021). Lastly, annotations of the network were propagated by the addition of *in silico* structure predictions by SIRIUS (confidence level 3). These different approaches resulted in a 16.5% annotation rate of the detected features (Table S1). The annotated features represented 12 different MFs in the network; thus, for 26 MFs no features were putatively annotated. Additionally, in some cases (e.g., MF 16, 18, and 19) it was not even possible for CANOPUS to predict the compound class based on the MS/MS spectra. Collectively, this demonstrates that the barley metabolome is highly complex and largely unknown.

CANOPUS (a part of the SIRIUS software) was applied for *de novo* chemical compound class predictions using the NPClassifier taxonomy (Kim et al., 2021). Overall, our results demonstrate that barley synthesizes a broad range of specialized metabolite classes, including a diverse array of terpenoids, phenylpropanoids, shikimates and alkaloids (Figure 2B). Terpenoids (MF 4) were largely unaffected by *P. teres* infection, with the exception of a putatively identified glucosylated conjugate of β-ionone (feature ID 3321; *m/z* 373.222 [M+H]^+^) which was significantly induced at 7 dpi (*p* = 0.001). In contrast, the production of many different phenylpropanoids, alkaloids and shikimates (i.e. MF 2 and 3) were induced during pathogen infection (Figure 2C).

Known barley metabolites were observed in the current study, with varying responses detected after infection. For example, all five hydroxynitrile glucosides (HNGs; epiheterodendrin, epidermin, sutherlandin, osmaronin, and dihydroosmaronin; MF 1), as well as the phenethylamine alkaloids hordenine and N-methyltyramine (MF 3) were detected, but did not significantly change in concentration in response to pathogen infection (Figure S2). In contrast, L-pipecolic acid (MF 3), *N*-hydroxy pipecolic acid glucoside (NHP-Glc; MF 26) and several methoxychalcones (MF 12) were only detected in the *P. teres* treated barley leaves. In MF 12, three methoxychalcone and flavanone features were annotated by spectral library matching as: 3,4,2’,4’,6’-pentamethoxychalcone, naringenin 5,7-dimethyl ether, and 2’,4-dihydroxy-3,4’,6’-trimethoxychalcone. Based on these annotations, we were able to further identify two neighboring nodes as 2’-hydroxy-3,4,4’,6’-tetramethoxychalcone and 4,6,2’’,4’’-tetramethoxychalcone 2’’-beta-glucoside at confidence level 2 and 3 by comparison to spectral information reported by Ube *et al*. (2021) and *in silico* structure prediction by SIRIUS, respectively (Table S1).

### Production of tryptophan-derived specialized metabolites are strongly induced in barley after *P. teres* infection

Overall assessment of the metabolic profile showed a significant alteration in metabolism from 4 dpi as a response to infection by *P. teres* as well as a variation caused by ontogenetic development (Figure S3). An assessment of LC-MS base peak chromatograms identified several peaks significantly induced after infection (Figure S4). In terms of absolute abundance, several tryptophan-derived specialized metabolites – particularly, tryptamine (*m/z* 161.10 [M+H]^+^ and *m/z* 144.08 [M-NH_3_+H]^+^), and three apparent hydroxylated tryptamine derivatives (*m/z* 177.10 [M+H]^+^ and *m/z* 160.08 [M-NH_3_+H]^+^) – were among the highest induced metabolites detected in our experimental setup. Mean tryptamine levels increased from being below the level of detection at 0 dpi to 2970 (± S.E. 915) nmol/g FW at 7 dpi, while the three different hydroxy tryptamine isomers were strongly induced from 0.28 (± S.E. 0.02), 0.34 (± S.E. 0.03), and non-detectable at 0 dpi to means of 137 (± S.E. 30), 359 (± S.E. 154) and 475 (± S.E. 104) nmol/g FW during *P. teres* infection, respectively (Figure 2D; Figure S4).

To determine the identity of the hydroxylated tryptamine derivatives, the acquired LC-MS data was mined in more detail. The three features all exhibited near identical fragmentation patterns but eluted at distinct retention times (Figure 3), leading to the hypothesis of different positional isomers of hydroxytryptamine. Authentic standards for serotonin, 6-hydroxytryptamine, *N*-hydroxytryptamine, 3-(2-Aminoethyl)-1H-indol-1-ol and 2-oxo-tryptamine (2OT) were obtained and/or chemically synthesized, whereby the chemical synthesis of 3-(2-Aminoethyl)-1H-indol-1-ol is reported for the first time. Furthermore, 4-hydroxytryptamine was obtained by transiently expressing PsiH; a characterized cytochrome P450 (Fricke et al., 2017) involved in biosynthesizing psilocybin in the fungi *Psilocybe cubensis* in *Nicotiana benthamiana,* and co-feeding with tryptamine. Peak 1 from the infected sample co-eluted with authentic serotonin (Figure 3), supporting previous findings that serotonin production is induced after pathogen infection in barley (Ishihara et al., 2017). Authentic 2OT co-eluted with peak 3 in the infected barley leaves (Figure 3). To further confirm the structure of this novel metabolite, peak 3 was purified and analysed by nuclear magnetic resonance (NMR) spectroscopy (Figure S5) and represents the first report of this metabolite from a biological system. The tested standards did not co-elute at the retention time of the remaining serotonin isomer (i.e. peak [2]) and in the experimental setup used, molecular ions of the two *N*-hydroxylated tryptamine derivatives [*N*-hydroxytryptamine and 3-(2-Aminoethyl)-1H-indol-1-ol] were not detected with our LC-MS method, and likely degraded. Authentic standards for the remaining α, β, 3’ and 7’ tryptamine hydroxylation positions were either unavailable or restricted.

**Figure 3:**
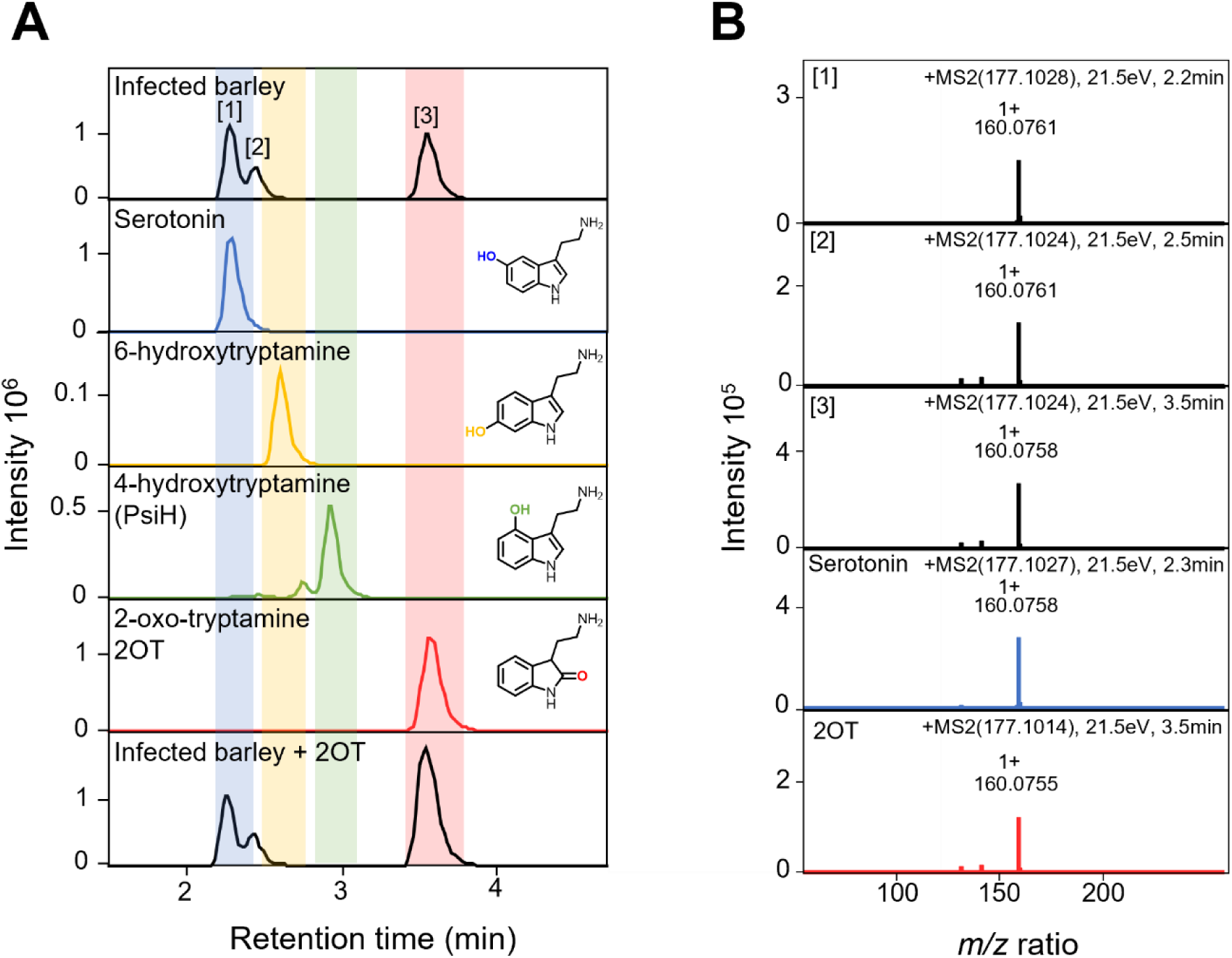
Production of serotonin and 2-oxo-tryptamine in barley leaves after fungal infection is confirmed by authentic standards. **(A)** Extracted ion chromatograms (EICs) (177.10; 160.08) matching the dominant ions of hydroxylated tryptamine. Top panel is a methanol extract of barley leaves infected with *P. teres* 7 days post inoculation (dpi). Panel 2, 3 and 5 are authentic standards of positional isoforms of hydroxylated tryptamine derivatives: serotonin (5-hydroxytryptamine; blue), 6-hydroxytryptamine (yellow) and 2-oxo-tryptamine (2OT; red), the spontaneous (oxidation product) of 2-hydroxytryptamine. Panel 4 is a methanol extract from *N. benthamiana* leaves expressing the 4-hydroxytryptamine (green) producing Cytochrome P450 (PsiH), fed with tryptamine 4 days post infiltration. Panel 6 is the extract from panel 1 spiked with 2OT. **(B)** MS2 fragmentation pattern for the parent ion (*m/z* 177.10 [M+H]^+^) of hydroxylated tryptamine isomers for peak [1], [2] and [3] and the authentic standards of serotonin (blue) and 2OT (red).

### Localization of hydroxylated tryptamine derivatives and other induced metabolites by MALDI-MSI

To investigate where 2OT and serotonin accumulate, Matrix Assisted Laser Desorption/Ionisation Mass Spectrometry Imaging (MALDI MSI) was performed on control and *P. teres* infected leaves, sampled at 4 dpi (n = 3). Ions corresponding to tryptamine (*m/z* 161.1073 [M+H]^+^; 144.0808 [M-NH_3_+H]^+^), 2OT and serotonin (*m/z* 177.1025 [M+H]^+^; 160.0757 [M-NH_3_+H]^+^) were detected and confirmed by independently assessing authentic standards in co-crystallization analysis. In an attempt to differentiate between serotonin and 2OT, tims (trapped ion mobility separation) analysis was performed, but no clear differentiation between the serotonin and 2OT ions was seen due to a low signal intensity observed for 2OT using this method (*data not shown*). In control samples, tryptamine was found to localize to mesophyll tissue, while trace levels of the ions corresponding to 2OT and serotonin were detected in the epidermal layer. After infection, ions corresponding to tryptamine and hydroxylated tryptamine derivatives are colocalized at the site of infection and found in the mesophyll tissue regions showing necrotic lesions and discoloration (Figure 4; Figure S7). However, within the infected tissue, tryptamine and hydroxylated tryptamine derivatives show different localization in the middle and outer regions, respectively.

**Figure 4:**
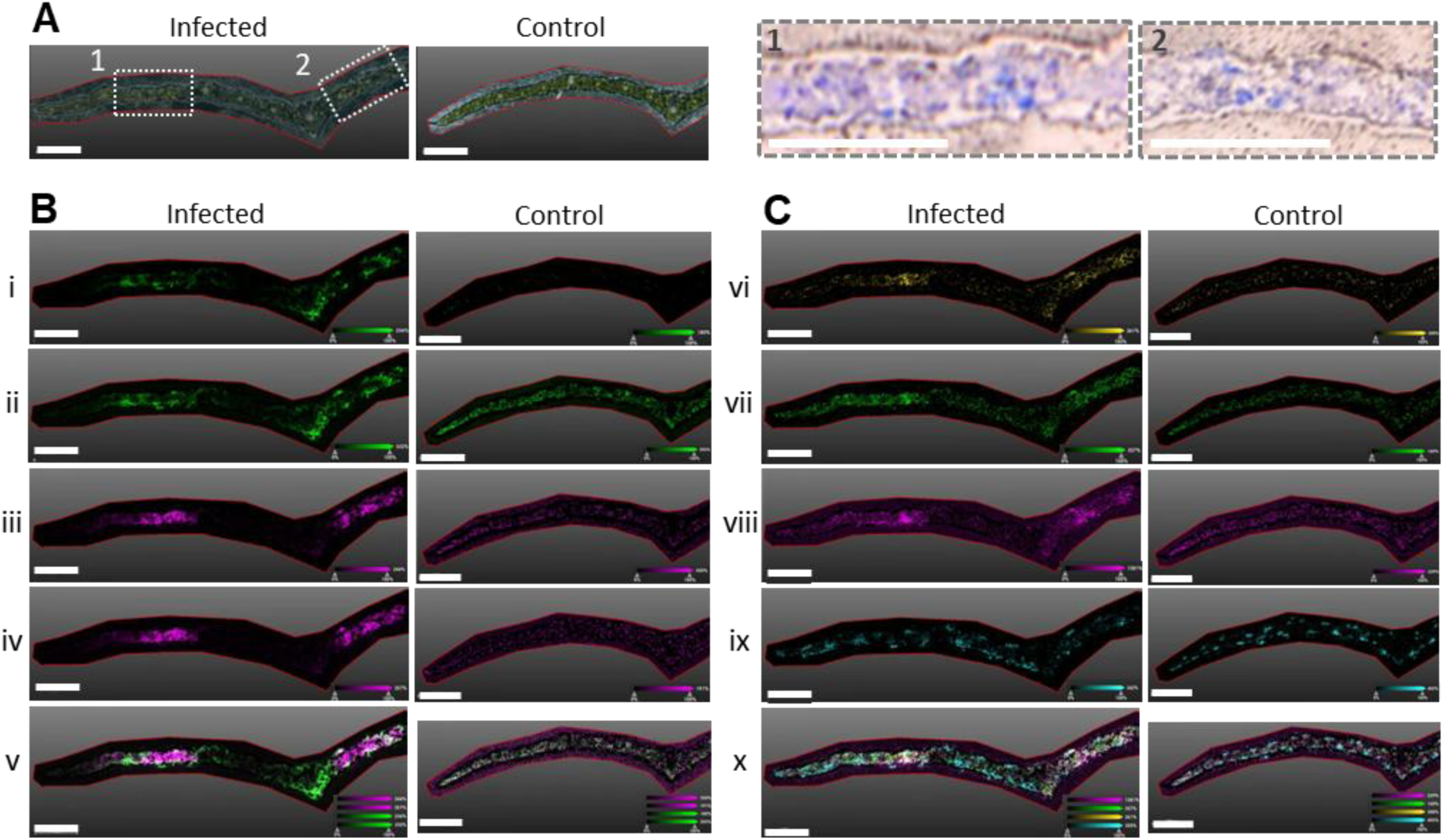
MALDI-MSI localization of selected metabolites in barley leaf cross sections after *P. teres* infection at 4 dpi with a spatial resolution of 20 µm. **(A)** Scanned images of representative infected and control leaves, with yellow necrotic tissue highlighted in the two dotted boxes. On the right, corresponding regions are color inverted with increased magnification, and necrotic tissue is indicated in blue. **(B)** Localization of tryptamine and hydroxylated tryptamine derivatives. Tryptamine (green) signals correspond to (**i**), *m/z* 161.1073 [M+H]^+^ and (**ii**) *m/z* 144.0808 [M-NH_3_+H]^+^ ions. Hydroxylated tryptamine derivatives (magenta) are detected as isobaric signals corresponding to (**iii**) *m/z* 177.1025 [M+H]^+^ and (**iv**) 160.0757 [M-NH_3_+H]^+^ ions. Overlay (**v**) of tryptamine and hydroxylated tryptamine derivatives show localization in middle and outer region of infected tissue, respectively. **(C)** Selected barley phytoalexins and phytoanticipins. The phytoalexins (**vi**) *N-*feruloylserotonin (yellow; *m/z* 391.1049 [M+K]^+^) and (**vii**) an unknown compound (green; compound ID 2191, *m/z* 132.0450 [M+H]^+^) are localized to the infected tissue regions. Signals corresponding to phytoanticipins (**viii**) hordenine (pink; m/z 204.0785 [M+K],) and (**ix**) sutherlandin (cyan; *m/z* 276.1077 [M+H]^+^) are detected in both control and infected leaf samples. Hordenine and sutherlandin are mainly detected in mesophyll and epidermal regions, respectively, although hordenine has a higher ion intensity at the sites of infection, as shown in the overlay image (**x**). Scale bar corresponds to 800 µm and 900 µm in infected and control samples, respectively. Color bars show intensity of detected ions. All images are generated with ± 5mDa error interval.

To identify and investigate the localization of other key metabolites induced after *P. teres* infection, random forest analysis was performed to rank the most important metabolite variables based on their contribution to the model accuracy at 4 dpi (Figure S6; Table S2). Within the top 15 most important contributing metabolites, three annotated metabolites were identified corresponding to 3,4,2’,4’,6’-Pentamethoxychalcone, *N*-feruloylserotonin, and tryptamine (MF 12, 2, and 3, respectively). MALDI-MSI analysis showed that two of these metabolites, *N-*feruloylserotonin (*m/z* 391.1049 [M+K]^+^) and an unknown compound (feature ID 2191; *m/z* 132.0450 [M+H]^+^), are also produced at the site of infection (Figure S8). In contrast, the phytoanticipins and apparently uninduced metabolites hordenine and the hydroxynitrile glucoside sutherlandin show an even distribution in the mesophyll and in clustered epidermal cells, respectively (Figure 4). Interestingly, an increased ion intensity of hordenine was detected at the infection sites. As overall hordenine concentrations did not differ between control and infected leaves (P = 0.57), this localization pattern may suggest metabolite relocation. Three other hydroxynitrile glucosides (dihydroosmaronin, epiheterodendrin, and epidermin; *m/z* 300.0844 [M+K]^+^) share the same localization pattern as sutherlandin. Osmaronin (*m/z* 298.0687 [M+K]^+^), however, shows a different localization pattern where the corresponding signal of its potassium adduct is also present in the mesophyll tissue of control and infected leaves (Figure S9). MALDI-MSI analysis also indicates a fine-tuned temporal and spatial response, as a region of tissue around the main midrib was found to accumulate tryptamine and the two phytoalexins prior to obvious necrotic lesions, and without co-localization of the biochemically downstream hydroxylated tryptamine derivatives. The other imaged biological replicates did not show accumulation of tryptamine and other metabolites at the midrib (or within vascular bundles; Figure S7, S8, and S10), so this accumulation does not suggest a localization associated with transport.

### Characterization of 2OTS and CYP71P10 (T5H)

#### Selection of candidate tryptamine hydroxylases

To identify potential tryptamine oxidases, a stepwise approach was used. First, transcriptome analysis was performed on RNA extracted from barley leaves infected with *P. teres* sampled at 0, 1, 2, 4 and 7 dpi. Water-treated leaves were included as a mock negative control. Given that a significant increase in 2OT and tryptamine was established at 4 and 7 dpi, respectively (Figure 2C), candidate monooxygenases were selected based on a log2 fold change cutoff of 2, late in the infection (i.e. 4 and 7 dpi). Specifically, CYPs and FMOs were targeted, previously shown to hydroxylate the indole backbone (Florean et al., 2025; Frey et al., 1997). This analysis resulted in 76 CYPs and six FMOs identified as potential candidates. To narrow the candidate list, a co-expression network using known genes for tryptamine biosynthesis was generated using PlantNexus (Zhou et al., 2022; Figure 5A&B), and assessed for the presence of the identified co-expressed monooxygenase enzymes using a mutual rank cutoff of 50. This resulted in the identification of three CYPs (CYP711A69; *HORVU4Hr1G079620*, CYP71U4; *HORVU7Hr1G043540* and CYP99A68; *HORVU2Hr1G004610*) and a Class B FMO from the N-

**Figure 5:**
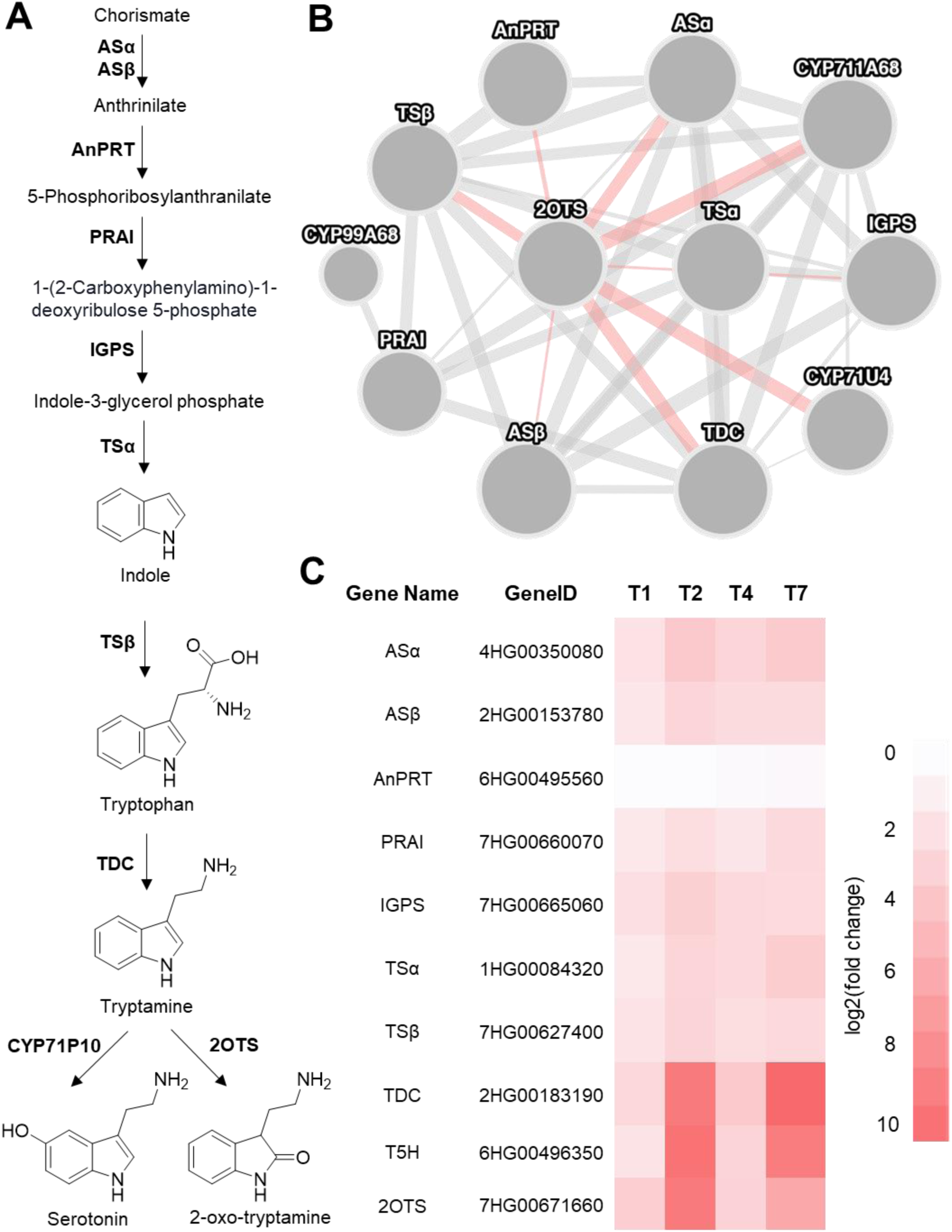
Co-expression analysis of genes involved in biosynthesis of tryptophan and tryptophan-derived defense metabolites. **(A)** Biochemical pathway for tryptamine, serotonin, and 2OT biosynthesis. ASα &Asβ (anthranilate synthase alpha/beta subunit), AnPRT (Anthranilate phosphoribosyltransferase), PRAI (Phosphoribosylanthranilate isomerase), IGPS (indole-3-glycerol-phosphate synthase), TSα & TSβ (Tryptophan synthase alpha/beta subunit), TDC (tryptophan decarboxylase), 2OTS (2-oxo-tryptamine synthase). **(B)** Tryptophan/tryptamine biosynthesis genes co-expression network generated in PlantNexus (Zhou et al., 2022). Mutual rank cutoff at ≤ 50. Red lines highlight the genes directly co-expressed with the FMO responsible for 2OT production (2OTS). **(C)** Log2(fold change) of tryptophan biosynthetic gene expression in barley plants infected with *P. teres* vs mock control at 1, 2, 4 and 7 dpi. GeneID is prefaced with ‘HORVU.RGT_PLANET.PROJ.’ for all genes.

OX3 family (HORVU7Hr1G121260) (Figure 5; Jiang et al., 2025). The identified FMO was expressed as two isoforms, differing by only two amino acids (Met and Gly) at the start codon, resulting in a slightly longer N-terminus. Accordingly, the isoforms were treated as the same gene for further characterization. An identified CYP71P10 exhibited a log2 fold change of approximately 10 at 2 dpi, and expression followed serotonin accumulation (Figure 2D; Figure 6). A corresponding barley HORVU gene identifier for this CYP71P10 does not exist in the first annotation of the barley *cv.* Morex genome, and therefore co-expression could not be assessed using PlantNexus. However, as the identified barley CYP71P10 is orthologous to the previously characterized tryptamine 5-hydroxylase CYP71P from rice (Fujiwara et al., 2010), this gene was included to assess its ability to hydroxylate tryptamine to form serotonin. Finally, candidate selection was expanded to include an orthologous gene approach, with two selected pathogen induced members of the CYP71C family included for further characterization: CYP71C115 (HORVU6Hr1G024850) and CYP71C34 (HORVU3Hr1G011870), despite having a co-expression mutual rank > 50. The CYP71C family has previously been shown to hydroxylate indole for benzoxazinoid biosynthesis in other grasses (Frey et al., 1997, 2009).

**Figure 6:**
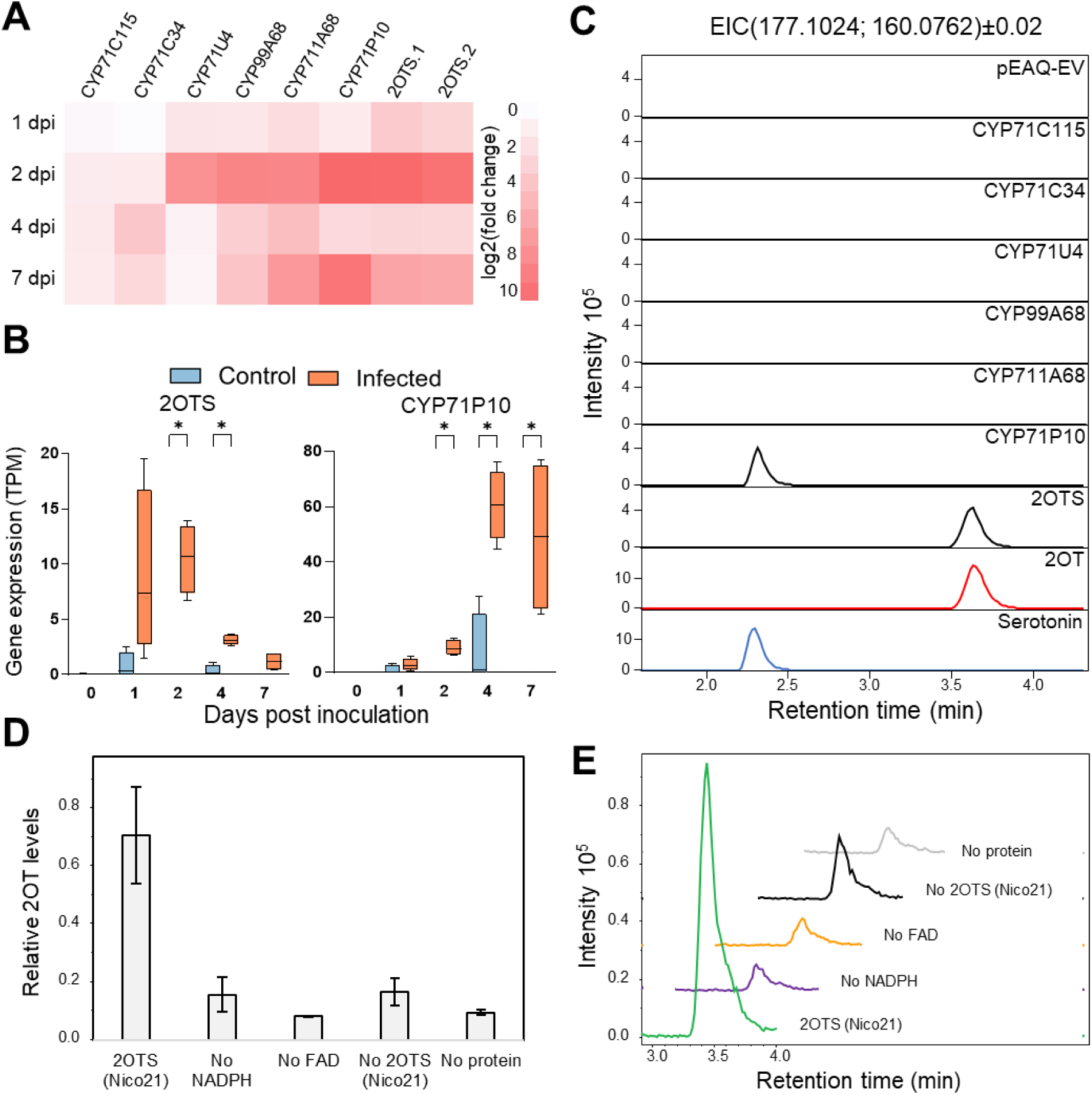
Characterization of hydroxylated tryptamine derivatives in *N. benthamiana* and *E. coli.* **(A)** Gene expression heatmap of log2 transformed fold change between infected and control at four different time points, 1 day post infection (dpi), 2 dpi, 4 dpi and 7dpi. **(B)** Transcript Per Million (TPM) value gene expression profile of two selected gene candidates, CYP71P10 (T5H) and 2OTS at 0 dpi, 1 dpi, 2 dpi 4 dpi and 7 dpi in both control and infected samples (n=4), asterisks denote significance by unpaired t-test at p-value < 0.05. **(C)** Extracted ion chromatograms (EICs) for the dominant ions of hydroxylated tryptamine-derived metabolites *m/z* 177.1024 [M+H]^+^ and *m/z* 160.0762 [M-NH_3_+H]^+^ from methanol extracts of *N. benthamiana* plants expressing 7 different monooxygenase candidates fed with tryptamine 4 days post infiltration and authentic standards for 2-oxo-tryptamine (2OT; red) and serotonin (blue). **(D)** Mean relative 2OT levels (± S.E.) measured in an *in vitro* activity assay of recombinant 2OTS supplied with tryptamine and cofactors FAD and NADPH. Background activity was measured using a crude lysate from the untransformed NiCo21 cells. **(E)** Representative EICs of the dominant ions of 2OT (*m/z* 177.1024 [M+H]^+^ and *m/z* 160.0762 [M-NH_3_+H]^+^) from the recombinant enzymatic assay.

#### An FMO from the N-OX3 family catalyses the hydroxylation of tryptamine to 2-oxo-tryptamine (2OT)

To test the activity of the seven candidate genes, *Agrobacterium tumefaciens* mediated transient expression was performed in *N. benthamiana* leaves. Leaves were supplied with exogenous tryptamine 4 d after infiltration followed by a methanolic extraction of metabolites 5 d after infiltration. LC-MS analyses of the extracts revealed that only two monooxygenase candidates were able to hydroxylate tryptamine: CYP71P10 and an FMO (Figure 6C).

Extracts from *N. benthamiana* plants expressing the CYP71P10 contained a feature matching the *m/z* and retention time of serotonin (Figure 6C), and we therefore conclude that the barley CYP71P10 is a tryptamine-5-hydroxylase (T5H). The metabolic extracts from plants expressing the FMO (HORVU7Hr1G121260) contained a feature matching *m/z*, retention time and fragmentation pattern of 2OT (Figure 6C). Heterologous expression of this FMO in *E. coli*, combined with *in vitro* crude enzymatic assays, demonstrated increased production of 2OT compared to controls, corroborating the *in vivo* experiment (Figure 6D-E). We therefore denote the identified FMO as a 2-oxo tryptamine synthase (2OTS).

2OTS possesses canonical class B FMO features: two paired Rossmann folds (G*X*G*XX*G) (CATH 3.50.50.60) binding FAD and NADPH and the FMO identifying motif (FXGXXXHXXXY/F) (Thodberg & Jakobsen Neilson, 2020). A BLAST search against a phylogenetic tree of the FMO N-OX family (Jiang et al., 2025) revealed 2OTS to be part of the N-OX3 family. An analysis of 2OTS in RGT Planet showed that RGT Planet possesses seven gene copies (> 98% identity) that are positioned in a tandem repeat gene cluster at the far end of chromosome 7 (Table S3). To assess if there are any natural *Hordeum* accessions that lack this FMO, we identified 2OTS gene orthologues across the pan-genome (76 accessions, including cultivated, landrace and wild *Hordeum* members; (Jayakodi et al., 2024). Genes with > 98% amino acid sequence identity to 2OTS were considered gene copies. All accessions within the pan-genome possess multiple copies of 2OTS, with gene copy numbers ranging from 2 to 8, all positioned at the end of pseudochromosome 7 in their respective genome assemblies (Table S3). Given that several of the tested monooxygenase genes are co-expressed and induced in barley leaves (Figure 5B; Figure 6A), it was hypothesized they may work together in concert to produce additional, novel metabolites. However, no additional products were observed when 2OTS, CYP71C115, CYP71C34, CYP71U4, CYP711A68 and CYP99A68 were transiently expressed together, with and without the addition of tryptamine (*data not shown*).

### 2-oxo-tryptamine is produced as a general defense response in *H. vulgare*

To assess whether serotonin and 2OT production represent a common defense response in barley, further biotic stress studies were performed using additional pathogens and cultivars. Specifically, *P. teres* (a hemibiotrophic pathogen) was also applied to barley *cv*. Golden Promise*, cv*. RGT Planet was infected with the bacterium *Pseudomonas syringae* (also hemibiotroph), and *cv*. Pallas was infected with the fungus *Blumeria hordei* (powdery mildew; biotroph). Distinct disease symptoms characteristic of the respective pathogens were observed in all cultivar-pathogen combinations (Figure 7), verifying effective infection. In all cases serotonin, 2OT and the novel unknown hydroxylated tryptamine derivative [2] were produced (Figure 7; Table S4). Notably, 2OT was consistently induced to relatively high levels across all treatments (Figure 7; Table S4). Interestingly, the typically susceptible *cv*. Golden Promise (Alhashel et al., 2021) exhibited minimal disease related symptoms after *P. teres* infection, and throughout disease development, the absolute levels of serotonin, hydroxylated tryptamine derivative [2], and 2OT were consistently lower than what was measured in the more affected *cv.* RGT Planet (Table S4). For instance, at 4 dpi, the levels were approximately 2.5, 3, and 8.5 times lower, respectively. Ratios between the three hydroxylated tryptamine isomers varied across the experiments. Generally, more serotonin and the novel hydroxylated tryptamine derivative [2] were produced relative to 2OT levels during *P. teres* infection compared to barley leaves treated with *P. syringae* and *B. hordei*. This may suggest that production of the different hydroxylated tryptamine isomers is differentially regulated depending on the specific pathosystem. Overall, these findings demonstrate that the induction of 2OT and related metabolites occurs broadly in barley-pathogen interactions, albeit with variation in their relative abundance.

**Figure 7:**
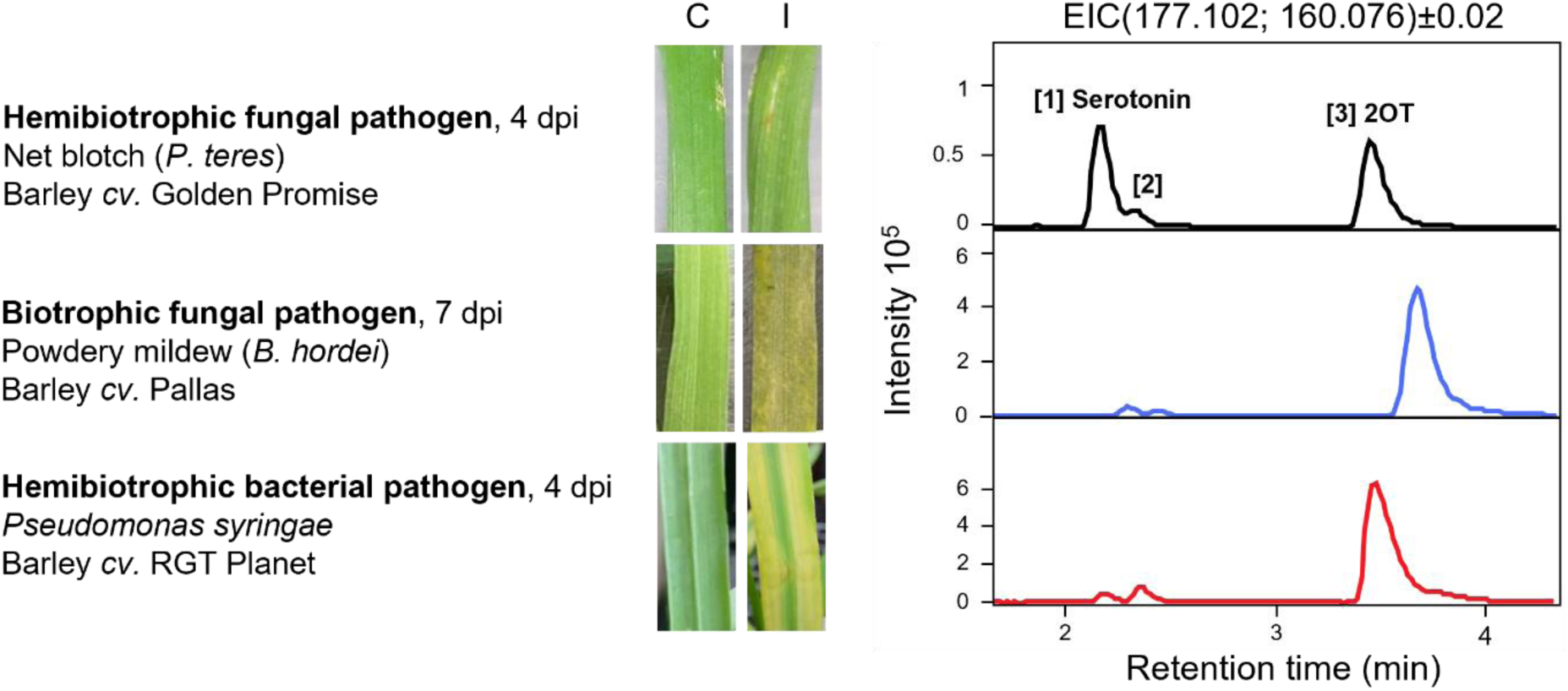
Induction of hydroxylated tryptamine derivatives in representative barley leaves of three different cultivars in response to infection by a biotrophic, and two different hemibiotrophic plant pathogens. An induced production of [1] serotonin, [2] unknown hydroxylated tryptamine derivative, and [3] 2-oxo-tryptamine (2OT) were observed in barley leaf tissue from *cv.* Golden Promise infected with *P. teres* (black), *cv.* Pallas infected with powdery mildew (*B. hordei*) (blue), and *cv.* RGT Planet infected with *P. syringae* (red). Representative picture of a control (C) and infected (I) leaf are shown.

### 2-oxo-tryptamine does not induce a systemic metabolic response

To investigate whether the production of 2OT and other tryptamine isomers triggers a distal signaling or mobile response known as systemic acquired resistance (SAR), barley (*cv.* RGT Planet) leaf tips were infiltrated with *P. syringae*. Methanol extracts sampled from both the tip (site of infection) and the base of the leaf at 7 dpi were subjected to LC-MS/MS analysis (Figure 8A). As an indicator of SAR, *N*-hydroxy pipecolic acid glucoside (NHP-Glc; (Bauer et al., 2021) was detected at the infection site but absent at the leaf base (Figure 8B). Some induction of NHP-Glc was also observed in mock-treated leaves, suggesting a wound response. As expected, at the site of infection (i.e. the tip), the analyses revealed an induction of serotonin, 2OT, and the unknown hydroxylated tryptamine derivative [2] (Figure 8C). However, no such induction was observed at the base of the leaf or in the mock-control leaves. Overall, no novel features were detected at the base of infected leaves compared to the base of mock-treated leaves, indicating a lack of SAR in this experimental setup (*data not shown*).

**Figure 8:**
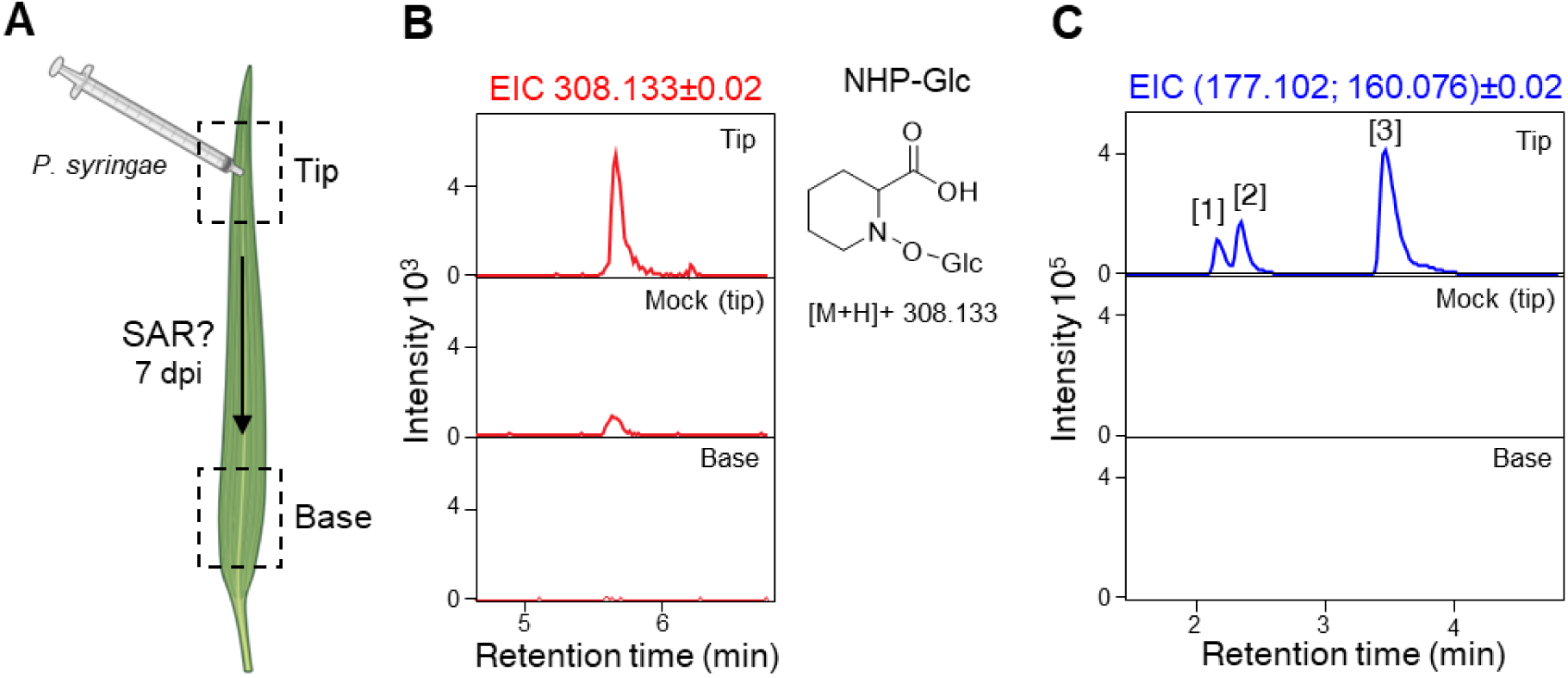
Investigation of a SAR induced metabolic response at the basal part of a barley leaf exposed to bacterial infection at the tip. **(A)** Schematic representation of the experimental setup (created using Biorender.com). Barley leaves were infected at the tip with *Pseudomonas syringae* pv. DC3000. Mock plants were infiltrated with water. 7 dpi, samples were collected from the site of infection (tip) and the base of the leaf for LC-MS/MS analysis. **(B) and (C)** EICs of *m/z* 308.133 [M+H]^+^ (±0.02) corresponding to NHP-Glc and the two dominant ions of hydroxylated tryptamine derivatives, *m/z* 177.10 [M+H]^+^ and *m/z* 160.08 [M-NH_3_+H]^+^ (±0.02). Abbreviations: SAR, systemic acquired resistance; EIC, extracted ion chromatogram; NHP-Glc, *N*-hydroxy pipecolic acid glucoside.

### Endogenous metabolism of tryptamine, 2OT and serotonin

The potential antimicrobial activity of 2OT and serotonin against *P. teres* was investigated with a bioactivity assay where spores of *P. teres* were incubated with 1 or 5 mM concentrations of serotonin, 2OT, and hordenine, a known barley defense alkaloid with reported antimicrobial properties (Ishiai et al., 2016), for 24 hours before being plated. After five days, fungal growth was evaluated and in contrast to hordenine, neither 2OT nor serotonin showed any significant inhibitory effect under these conditions (Figure S11). Therefore, it was hypothesized 2OT and serotonin might be intermediates and further metabolized in barley. To investigate this hypothesis, barley leaves were supplied with exogenous tryptamine, 2OT and serotonin. LC-MS analysis was performed on extractions from leaves infiltrated with either 100 µM tryptamine, 2OT, serotonin, or water (control) 24 hours post infiltration. Leaves fed with compound were used to individually compare and calculate fold change and p-value against control using MS-DIAL (Tsugawa et al., 2015). Features with a fold change (Max/Min) value above 2 and a p-value below 0.05 were manually identified if possible and cross-referenced against the molecular network of infected barley leaves (Fig. 2). Barley leaves infiltrated with exogenous tryptamine produced two features with *m/z* 177.10

[M+H]^+^ and retention times (rt) corresponding to previously observed hydroxylated tryptamine isomers: an unidentified hydroxylated tryptamine isomer at rt 2.3 and 2OT at rt 3.4 (Table 1). Notably, no increase in serotonin levels were detected. Additionally, two other features, *m/z* 193.10 and *m/z* 705.19, showed fold increases of 123 and 3, respectively. The feature with *m/z* 193.10 matched the theoretical mass of dihydroxytryptamine but did not align with the retention time of an authentic standard for 3-(2-Aminoethyl)-3-hydroxyindolin-2-one (AEHI), a phytoalexin reported by (Ishihara et al., 2017). The feature with *m/z* 705.19 was not present in the molecular network, nor was its structure elucidated.

**Table 1:** Metabolite features produced in barley leaves supplied with tryptamine, 2OT and serotonin. If a feature was also identified in the barley metabolomics network (Figure 2), the feature ID from the molecular network is noted. Abbreviations: 2OT, 2-oxo-tryptamine; AEHI, 3-(2-Aminoethyl)-3-hydroxy-indolin-2-one; RT, retention time.

| Exogenous compound | *m/z* | RT (min) | ANOVA P-value | Fold Change (Max/Min) | Manual Annotation | Feature ID | Notes |
| --- | --- | --- | --- | --- | --- | --- | --- |
| Tryptamine | 144.0815 | 5.97 | 9.1E-05 | 45 | Tryptamine | 2040 |  |
|  | 161.1081 | 5.95 | 1.2E-04 | 36 |  | 2026 |  |
|  | 160.0772 | 2.32 | 1.2E-02 | 33 | Hydroxylated tryptamine derivative [2] | 1551 |  |
|  | 177.1032 | 2.32 | 1.9E-02 | 55 |  | 1550 |  |
|  | 177.1029 | 3.41 | 4.2E-02 | 5 | 2OT | 1763 |  |
|  | 193.0979 | 3.60 | 3.9E-02 | 123 | Unknown | 1791 | *m/z* matches dihydroxytryptamine |
|  | 705.1865 | 1.14 | 2.0E-03 | 3 | Unknown | N/A |  |
| 2OT | 160.0761 | 3.41 | 7.2E-04 | 97 | 2OT | 1759 |  |
|  | 177.1029 | 3.41 | 9.0E-04 | 103 |  | 1763 |  |
|  | 146.0608 | 1.59 | 3.0E-02 | 19 | AEHI | 1279 | (Ishihara et al., 2017) |
|  | 193.0974 | 1.59 | 3.1E-02 | 50 |  | 1276 |  |
| Serotonin | 160.0765 | 2.13 | 5.4E-04 | 556 | Serotonin | 1480 |  |
|  | 177.1031 | 2.13 | 7.9E-05 | 294 |  | 1484 |  |
|  | 452.2035 | 2.13 | 6.7E-05 | 670 | Unknown | 1486 |  |
| | 191.0822 | 2.15 | 1.2E-03 | 27 | Unknown | 1493 | *m/z* matches $\alpha/\beta$-oxo-serotonin |
|  | 221.0930 | 2.14 | 1.3E-03 | 28 | Unknown | 1485 |  |

In leaves infiltrated with 2OT, only 2OT itself and a feature corresponding to the *m/z* and retention time of AEHI were observed, demonstrating that barley can metabolize 2OT into AEHI. In extracts from leaves infiltrated with serotonin, three features (besides serotonin), *m/z* 452.20, *m/z* 191.08, and *m/z* 221.09, showed increased levels (Table 1). While the structures of these features were not elucidated, their presence in the molecular network supports the hypothesis that barley produces these compounds in response to *P. teres* infection.

## Discussion

### Barley induces a strong metabolomic defense response to *P. teres* infection

The application of untargeted metabolomics is a pivotal tool within plant science research to enhance the understanding of metabolic reconfiguration in response to various stressors, thereby contributing to improved crop resilience and yield (Bozza et al., 2024). A major challenge within untargeted LC-MS based metabolomics relates to compound identification due to the vast amount of unknown compounds (Giera et al., 2022; Muñoz-Hoyos & Stam, 2023). Here we used feature-based molecular networking (FBMN), which is a part of the GNPS2 platform, for global visualization and chemical annotation of our generated mass spectrometry dataset. In molecular networking, experimental mass spectra from detected chromatographic features are aligned against one another and connected based on spectral similarity. Each feature is depicted as a node in the network, and related nodes are grouped in molecular families (MFs) (Nothias et al., 2020). FBMN can distinguish between isomers with very similar MS2 fragmentation spectra and was a useful tool to differentiate between three key hydroxylated tryptamine derivatives induced after pathogen infection (Figure 2). Additionally, by combining the generated FBMN with *in silico* annotation tools (e.g. SIRIUS), the network nodes can function as an anchor to annotate structurally similar metabolites in the same MF and help decode the large unknown chemical space.

Here, 16.5% of metabolites were annotated representing a relatively high annotation rate for an untargeted metabolomics dataset (De Jonge et al., 2022). A large proportion of barley features (83.5%), however, remain unannotated. Given that many unknown metabolites are strongly induced in infected samples (e.g. MF 13 and MF 15; Figure S12), much remains to be elucidated regarding the barley metabolic defense response. However, it should be noted that a part of the unannotated metabolic space might not represent unknown metabolites, but instead reflect in-source fragmentation (Giera et al., 2024). Additionally, biochemically very similar metabolites sometimes produce fragmentation patterns that are so different, they do not cluster in the same MF in the network thus adding another layer of complexity to the annotation process. In our study these challenges were exemplified by e.g. the parent ion for the phytoalexin AEHI (*m/z* 193.09 [M+H]^+^) being present as a singleton, while the in-source fragment ion *m/z* 146.06 for AEHI actually is connected to the identified hydroxylated tryptamine isomers in MF 3 (Figure 2B and C; Table S1). Moreover, the in-source fragment ion was initially wrongly annotated as isoquinolone. The correct annotation as AEHI was therefore only possible because of the availability of authentic standard.

The metabolomic pipeline employed in this study shows that barley produces a wide arsenal of specialized metabolites, including alkaloids, terpenoids, hydroxynitrile glucosides and phenylpropanoids (Figure 2). Some metabolites, such as the hydroxynitrile glucosides, hordatines, and hordenine, were unaffected by the biotic stress applied here while other annotated metabolite classes are induced. Specifically, indole alkaloids and in particular tryptophan-derived metabolites, hydroxycinnamic acid amides (HCAAs) (e.g. *p*-CHDA, feruloyltyramine, sinapoylagmatine, and *N*-feruloylserotonin), and phenylpropanoids were induced after fungal infection. The metabolites produced, and the metabolic shift in response to infection is in line with other studies (Hamany Djande et al., 2023; Ishihara et al., 2017; Lemcke et al., 2021; Ube et al., 2021; von Röpenack et al., 1998). For example, aromatic amino acid metabolism, phenylpropanoid and isoquinoline alkaloid biosynthesis are induced in the *P. teres* susceptible barley *cv.* Hessekwa (Hamany Djande et al., 2023). Recently, several barley-specific labdane-related diterpenoids, hordedanes, were identified as important phytoalexins in response to fungal infections (Y. Liu et al., 2024). These metabolites were observed by Liu and colleagues in root exudates, but were not identified in our leaf methanol extracts, indicating that the production of hordedanes is root specific.

### Tryptophan-derived metabolites play an important role in barley defense response

In regard to the hypothesized tryptophan-derived metabolites, tryptamine, and three hydroxylated tryptamine isomers were identified to be strongly induced after infection in multiple barley cultivars and in response to multiple plant pathogens with different pathogenic lifestyles (Figure 2D; Figure 7). Two of the hydroxylated tryptamine derivatives were confirmed to be serotonin and 2-oxo-tryptamine (2OT) (Figure 3). The identity of the third hydroxylated tryptamine isomer remains unknown. Ishihara *et al*. reported the induction of a di-hydroxylated tryptamine derivative at positions 2’ and 3’ (AEHI) in barley after pathogen infection, demonstrating that barley has the capacity to hydroxylate tryptamine at the 3’ carbon. This suggests that the unidentified hydroxylated tryptamine derivative could potentially be 3-hydroxytryptamine. Furthermore, the metabolic network identified many features with an indolic backbone, including 3-methyl-2-oxindole, indoline, *N*-methyltryptamine (Table S1; Figure 2). These data support the hypothesis that tryptophan is critically important for the execution of defense during barley infection. While this study assesses the non-volatile metabolome of barley, volatile organic compounds (VOCs), have also been shown to play an important role in how barley responds to pathogen infection. Nonanal and the β-ionone are emitted from barley leaves infected with *Pseudomonas syringae pv. japonica*, and are proposed to trigger a plant-to-plant SAR defense response (Brambilla et al., 2022). In line with this result, we putatively identified glucosylated conjugates of β-ionone and other related β-ionone-like terpenoids, which may be the storage form of β-ionone prior to emission. Based on the strong induction of the genes related to indolic compounds, it would be highly relevant to perform untargeted VOC analysis of barley and other cereals to assess whether indole-related metabolites are both emitted as well as stored. Indeed, volatile indole is an important VOC following herbivore attack in maize (Erb et al., 2015) and rice (Zhuang et al., 2012).

### Pathway elucidation of 2OT and serotonin in barley

Through a combination of heterologous expression in tobacco and *E. coli*, 2OTS was shown to catalyze the conversion of tryptamine to 2-oxo-tryptamine. It is possible that 2OTS hydroxylates tryptamine at the 2’ position, which then tautomerizes to 2OT, as reported for oxindole (Kaur et al., 2016). The identified 2OTS is a class B FMO from the so-called N-OX super family (Jiang et al., 2025 [Chapter VI]; Chapter III) one of several major FMO families in plants. Recent cross-kingdom evolutionary analysis of class B FMOs further classifies the N-OX superfamily as belonging to a diverse FMO clade (Clade IV) comprising almost exclusively plant species, but also contain an FMO from the protozoan *Trypanosoma cruzi* (Nicoll & Mascotti, 2023). Other plant FMOs families include the YUCCAs (Clade III) and S-OXs (Clade II), as well as a single Baeyer-Villiger monooxygenase (BVMO) reported from the moss *Physcomitrella patens* (Beneventi et al., 2013; Nicoll & Mascotti, 2023; Thodberg & Jakobsen Neilson, 2020). The N-OX family was historically named based on previously characterized FMOs shown to hydroxylate nitrogen in small molecules (Thodberg et al., 2020). This naming was proposed on the function of only two FMOs: FOS1 from ferns that performs two subsequent *N*-hydroxylations on phenylalanine leading to phenylacetaldoxime as part of cyanogenic glycoside biosynthesis (Thodberg et al., 2020) and Arabidopsis FMO1 that catalyzes a *N*-hydroxylation of pipecolic acid to form N-OH pipecolic acid eliciting a SAR signaling response (Chen et al., 2018; Hartmann et al., 2018). Heterologous expression in *Nicotiana benthamiana* has also demonstrated substrate promiscuity, such that *At*FMO1 can hydroxylate a second small molecule putatively identified as *N*-hydroxylate proline (Holmes et al., 2019). Since then, other N-OX FMOs have been characterized, shown to perform both *N*- and *C*-hydroxylations. A N-OX3 FMO was recently discovered in Darwin’s Orchid (*Angraecum sesquipedale*), and is responsible for dual *N-*hydroxylations of phenylalanine and leucine to form their respective aldoximes as pollinator-attracting volatile scent molecules (Jiang et al., 2025). The BX N-OXs from *Aphelandra squarrosa*, *Lamium galeobdolon* and *Consolida orientalis* perform a dual *C*-hydroxlation on the C2 and C3 positions of indole, as intermediates in benzoxazinoid biosynthesis (Florean et al., 2023, 2025). Finally, a FMO from *Polygonum tinctorium* capable of oxidizing indole to form indigo when expressed in *E. coli* (Inoue et al., 2021). These findings necessitate a reevaluation of the assumption that FMOs from the N-OX family primarily hydroxylate nitrogen atoms, and instead suggest that this family can perform both *C*- and *N*-hydroxylations on small aromatic and ring-like metabolites.

The biosynthetic pathway of serotonin in plants has been elucidated in rice, tomato and green foxtail all by enzymatic characterization of a CYP450 with tryptamine-5-hydroxylase activity (Fujiwara et al., 2010; Commisso et al., 2022; Dangol et al., 2022). A BLAST of *Hv*CYP71P10 identified in this study showed that it shared 87%, 80% and 55% sequence identity with OsCYP71P1 (rice), SvT5H (green foxtail) and SlT5H (tomato), respectively. Cytochrome P450 nomenclature dictates that enzymes sharing 55% identity belong to the same subfamily, placing all the genes in the CYP71P subfamily (Nelson, 2005). In barley, the function of CYP71P10 has previously been indirectly characterized by the genetic analyses of the *nec3* mutant, a disease lesion mimic mutant with impaired serotonin biosynthesis (Ameen et al., 2021). Confirmation of CYP71P10 enzymatic function via heterologous expression in yeast was unsuccessful, so the tryptamine-5-hydroxylase capability of CYP71P10 reported here opens avenues for further genomic studies using the *nec3* mutant.

### The role of serotonin

Here we show serotonin is significantly induced at 4 dpi, with levels increasing as the *P. teres* infection progressed. Serotonin has previously been found to be produced in response to pathogens in barley and other grasses (Ishihara et al., 2008, 2017; Lemcke et al., 2021; Powell et al., 2016). The defense-related roles attributed to serotonin in Poaceae are multifaceted. Our best understanding of the role of serotonin in barley is facilitated by the *nec3* mutant which develops atypically large necrotic lesions when exposed to a variety of penetrative pathogens. It has therefore been postulated that serotonin is a negative regulator of programmed cell death via an interaction with reactive oxygen species (ROS) and links to cuticle stability (Ameen et al., 2021). In rice, serotonin accumulates at sites of hypersensitive response (HR) lesions and is therefore proposed to help confine oxidative damage through its antioxidant activity, thereby preventing excessive cell death beyond infection sites (Hayashi et al., 2016).

A similar necrotic lesion phenotype is also observed in a rice mutant devoid of serotonin, known as the *sl* (Sekiguchi Lesion) mutant that lacks a functional *Os*CYP71P1 (Fujiwara et al., 2010). Rice *sl* mutants have enhanced pathogen susceptibility and are unable to deposit so-called ‘brown material’ (Ishihara et al., 2008). The ability to produce the brown material is restored after supplementation of serotonin and by supplementing infected *sl* mutants with radiolabeled serotonin, 80 percent was deposited into the cell wall. Taken together, these results suggest that serotonin could also function as a key metabolite in physical cell wall fortification, helping to block pathogen entry. MALDI results show serotonin to be mainly localized at the sites of infection, which supports this hypothesis. Interestingly, trace levels of serotonin were also detected in the epidermis of uninfected barley leaves and supports the findings within cuticle stability (Ameen et al 2021). The role of serotonin within biotic interactions in complex, such that plants may be more resistant to insect pests when serotonin biosynthesis is suppressed (Lu et al., 2018).

Possibly, serotonin could be involved in the recruitment of beneficial microbes during pest attack or infection. A recent study has shown that serotonin is a keystone metabolite exuded from switch grass roots during nutrient deficiency to recruit beneficial microbes (Baker et al., 2024). Given that the induced systemic resistance is activated during infection by a hemibiotrophic pathogen, serotonin may likely also be exuded from barley roots to facilitate microbial interactions in a similar way. Overall, serotonin is an important cross-kingdom metabolite, and further investigations using genetic models are highly relevant for uncovering its role in plant–microbe interactions and systemic defence signaling.

### The potential roles of 2OT

The role of 2OT is in its nature unknown as the metabolite has not previously been reported in plants. We investigated three different hypotheses regarding the role of 2OT. The hypotheses were: [1] 2OT is a signaling compound involved in systemic acquired resistance in barley, [2] 2OT is a phytoalexin induced after infection as a chemical defense response, [3] 2OT is an intermediate in the biosynthetic pathway of another defense compound.

[1] In an experiment where only the tip of barley leaves was infected, we analyzed whether the production of 2OT led to metabolic changes in the base of the leaf. We used the recognized SAR metabolite *N*-hydroxy pipecolic acid glucoside (NHP-Glc) as an indicator of signaling throughout the leaf. Regardless of the fact that we observed induction of 2OT and NHP-Glc in the tip of infected leaves no metabolic response was observed in the base of the leaves. In this experimental setup no signaling leading to a metabolic response was initiated by accumulation of 2OT, suggesting that the role of 2OT is not involved in signaling.

[2] To test the direct chemical defense capabilities of 2OT as a phytoalexin we performed a bioassay against *P. teres*. The pathogen was exposed to differing concentrations of 2OT, serotonin and hordenine. While hordenine exhibits clear inhibition of *P. teres* germination no such inhibition was observed for either 2OT or serotonin. The lack of antimicrobial activity of these hydroxylated tryptamine derivatives questions their role as direct phytoalexins. Given that 2OT is structurally similar to serotonin, and MALDI MSI shows these two compounds co-localize during infection, it is possible that 2OT may be incorporated into the fortification of cell walls and or the cuticle.

[3] To determine whether barley is able to further metabolize 2OT, we supplied healthy barley plants with exogenous 2OT through infiltration and analyzed changes in the metabolic profile compared to control plants. We observed a single compound increasing as a result of 2OT supplementation, the previously identified phytoalexin 3-(2-Aminoethyl)-3-hydroxyindolin-2-one (AEHI) in barley (Ishihara et al., 2017). This finding suggests that 2OT is part of the biosynthesis of AEHI as an intermediate, which is supported by the co-localization of AEHI at the site of infection (Figure S7). It is possible that 2OT may be an intermediate for other indolic compounds, but in this setup the downstream catalyzing enzymes may not be available i.e., they are only expressed when activated by specific defense gene cascades. For example, the detected serotonin-conjugates in our metabolic analysis were not measured in the feeding study.

## Conclusions

In this study, we investigated the metabolic defense response of barley following infection by the hemibiotrophic fungal pathogen *Pyrenophora teres* f. *teres*, the causal agent of net blotch. This disease poses a significant threat to global barley production, with yield losses of up to 40% reported (Clare et al., 2020). Based on previous pathogenic studies, we hypothesized that tryptophan-derived specialized metabolism would play a key role in the barley defense response. Our results support this hypothesis, revealing a strong activation of tryptophan metabolism after *P. teres* infection. In particular, we identified serotonin and two hydroxylated tryptamine isomers as key metabolites induced at the infection site. One of these, 2-oxo-tryptamine (2OT), represents a previously unreported plant metabolite. We further demonstrate that serotonin and 2OT are biosynthesized from tryptamine by CYP71P10 and a flavin monooxygenase (2OTS) from the N-OX3 subfamily, respectively. These findings highlight the importance of tryptophan-derived specialized metabolism in barley defense, potentially contributing to cell wall fortification and the production of bioactive phytoalexins. This work lays a foundation for further exploration of metabolic defense pathways and may inform strategies for enhancing disease resistance in cereal crops.

## Materials and Methods

### Barley plant material and cultivation

In this study, three 2-rowed spring barley (*Hordeum vulgare*) cultivars *cv.* Golden Promise, *cv.* RGT Planet, and *cv.* Pallas were used. Golden Promise is a semi-dwarf mutant resulting from gamma-ray treatment of *cv.* Maythorpe (Sigurbjornsson & Micke, 1969). Previously, it was extensively used in the malting and whisky industry, especially in Scotland. However, at present Golden Promise receives considerable interest due to its superior genetic transformability compared to other barley cultivars (Schreiber et al., 2020). Barley *cv.* RGT Planet is a modern 2-rowed spring barley cultivar developed for malting purposes. Pallas is an erectoides x-ray mutant of the barley *cv.* Bonus isolated in 1947 and released in 1958 by the Swedish Seed Association, Svalöf. It quickly became widely adopted in Western Europe due to its high yield, high nitrogen utilization, and resistance to lodging (Gustafsson & Ekman, 1967). In this study, we used the specific Pallas line P-02, which carries the *Mla3* powdery mildew resistance gene (Kølster et al., 1986). Seeds were imbibed on wet filter paper in Petri dishes overnight at 4 °C and subsequently germinated at approximately 1 cm depth in one row in soil-medium (Pindstrup substrate no. 2, Pindstrup Mosebrug A/S, Denmark). About 12 seeds were planted per pot (13 x 13 cm). Plants were grown in a growth chamber under controlled light and temperature conditions: cycles of 16h light/8h darkness with 200 µE m-2 s-1 light intensity (fluorescent tubes, Philips Master IL-D 36 w/865, France). Temperature and relative humidity were 19 °C/50-60 % in light and 16 °C/80-90 % in darkness, respectively.

### Pathogen inoculum preparation

#### Pyrenophora teres f. teres

The hemibiotrophic fungal pathogen *Pyrenophora teres f. teres* (isolate CP2189), was used. It causes the disease net blotch – a major foliar disease of barley (Tini et al., 2022). The pathogen was grown for two weeks on grass agar plates (filtrate of 32.5 g/l of boiled clover-rich grass fodder pills for cattle and 20 g/l agar) according to (Jørgensen et al., 1996). On the day of inoculation, conidia were washed off with sterile deionized water. Subsequently, spores were counted using a haemocytometer and the inoculum concentration adjusted to 3-5000 spores/ml according to Pandey et al. (2021).

#### Pseudomonas syringae pv. tomato

Cultures of *Pseudomonas syringae pv. tomato* DC3000 were provided by Dr. Kenneth Madriz Ordenana, University of Copenhagen, Denmark. They were grown at 28 °C in LB medium supplemented with rifampicin (25 µg/mL) until reaching late log phase. The bacteria were harvested by centrifugation (10 min at 2500 *x* g), washed once in 10 mM MgCl_2_ and resuspended in 10 mM MgCl_2_ and 0.01% (v/v) Silwet L-77 adjusted to an optical density (OD) at 600 nm = 0.2-0.3.

#### Blumeria hordei

The biotrophic pathogen barley powdery mildew fungus (*Blumeria hordei;* syn. *B. graminis f.sp. hordei* (Liu et al., 2021) isolate C15 (*AVR_A1_*) were kindly provided by Dr. Mads Eggert Nielsen. The fungal inoculum was propagated on the barley *cv.* Pallas line P-02 by weekly transfer.

### Pathogen inoculation and experimental design

Fungal infection studies (*Pyrenophora teres* and *Blumeria hordei*) were independently repeated twice, both confirming the production of 2OT, serotonin, and other tryptophan-derived compounds.

#### Pyrenophora teres f. teres

At the two leaf stage (BBCH 12;Lancashire et al., 1991), the second developed leaves were fixed in a horizontal position (adaxial side facing up) on bent Plexiglas plates using un-bleached cotton strings (Jørgensen et al., 1996). Plants were left to acclimatize in the new position for three days before inoculation. Subsequently, fixed leaves were spray inoculated with a spore suspension until run-off. Immediately after inoculation, plants were covered in plastic bags to ensure 100% RH and kept in darkness in the growth chamber. Control plants were sprayed with water and maintained under the same conditions as infected plants. Samples were collected at 0 (non-treated), 1, 2, 4 and 7 dpi. For each sample type (non-treated, infected and control) at each time point, five independent biological replications (individual fixed leaves) were cut into pieces and randomly divided in two Eppendorf tubes per leaf – one for metabolomic and one for transcriptomic analysis, respectively. Immediately, samples were snap-frozen in liquid N_2_ to quench metabolic activity and stored at −80 °C until metabolite or RNA extraction.

#### Pseudomonas syringae pv. tomato

The prepared bacterial suspension was infiltrated into the abaxial side of the second forming leaf by syringe injection as described previously (Katagiri et al., 2002) using a 1-mL needleless syringe. Post infiltration, plants were covered in plastic bags to ensure high humidity and kept in the growth chamber. Control plants were infiltrated with water and maintained under the same conditions as infected plants. For the systemic response experiment, leaves were only infiltrated at the leaf tip and sampled tissue was collected from the base of the leaf, 5 cm from the site of infiltration. Three individual biological replications were sampled on 4 and 7 dpi, snap-frozen and stored at −80 °C until metabolite extraction.

#### Blumeria hordei

One week old plants were fixed on Plexiglas plates as above and inoculated with *B. hordei* from sporulating *cv*. Pallas line P-02 plants as previously described (Jørgensen et al., 1996). Leaf material was collected from control and infected leaves (n = 3) at 4 dpi, snap frozen and stored at −80°C until metabolite extraction.

### Transient expression and substrate feeding in *Nicotiana benthamiana*

#### Cloning of Gateway constructs

All genes were commercially synthesized (Genscript) with an N-terminal 6xHIS tag and flanking attL sites in pUC57 (ThermoScientific) essentially functioning as entry vectors. Expression vectors were generated by gateway recombination with PEAQ-HT-DEST1 and gene containing entry vectors. PEAQ-HT (empty vector) was used as a control (Sainsbury et al., 2009). Expression clones were verified by whole plasmid sequencing (Plasmidsaurus, Azenta & UnveilBio).

#### Transformation of Agrobacterium tumefaciens and substrate feeding

Verified plasmids were transformed into electro-competent *Agrobacterium tumefaciens* strain AGL-1 (Lazo et al., 1991). Transformed agrobacteria were grown in YEP media O/N at 28°C. Cell were harvested by centrifugation (4000 *x* g, 10 min) and suspended in infiltration buffer, pH 5.6 (50 mM MES, 2 mM sodium phosphate, 0.01% Silvet L-77, 0.5% D-Sucrose, 200 µM acetosyringone) to OD_600_ = 0.8 and incubated at room temperature for (1-)3 hours. *Agrobacterium* suspensions were injected using a 1 mL syringe on the abaxial leaf surface of 4-5 weeks old *Nicotiana benthamiana* plants. Plants were placed in a greenhouse (21 °C day/19 °C night) for 4 days before 100 µM tryptamine in infiltration buffer was injected in the same way as the *Agrobacterium* suspension. After one day, three leaf discs from separate leaves from one plant were pooled with a 1 mm steel ball bearing and flash-frozen in liquid N_2_ and stored at −80 °C until metabolite extraction.

### Heterologous expression and crude enzyme preparation of 2OTS

The 2OTS gene from *Hordeum vulgare* (Phytozome Gene ID: HORVU7Hr1G121260) was codon-optimized and cloned into the pNIC28-Bsa4 vector (Savitsky et al., 2010) carrying the coding sequence for a TEV-cleavable N-terminal His-tag and kanamycin resistance, and transformed into NiCo21 pRARE2 (DE3) *E. coli*. A single colony was picked from a fresh LB-agar plate (containing 50 µg/mL kanamycin and 25 µg/mL chloramphenicol) and the seed culture for 2OTS protein expression (MW: 60.77 kDa) was grown overnight (37 °C, 250 rpm) in 50 mL rich TB medium supplemented with kanamycin and chloramphenicol. Furthermore, a 2 L 2OTS expression batch was initiated by inoculating each litre of TB medium (supplemented with kanamycin and chloramphenicol) with 20 mL bacterial seed culture in 5 L baffled flasks. Pre-induction cultures were incubated (37 °C, 250 rpm) until OD₆₀₀ = 1.2–1.5. Subsequently, protein expression was induced with 0.5 mM IPTG, the culture was incubated O/N at 18 °C (150 rpm, ∼16–18 hours) and the *E. coli* cells were harvested by centrifuging in a SORVALL LYNX 6000 centrifuge (4 °C, 5000 × *g* for 15 min). *E. coli* cells were resuspended in lysis buffer (100 mM Tris pH 7.5, 300 mM NaCl, 1 mM MgCl₂, 0.5 mM TCEP, 0.04 mg/mL lysozyme, protease inhibitor (1 tablet/50 mL buffer), Benzonase (Sigma-Aldrich), and NP-40 (0.05%)) (5 mL/g cells) and incubated on ice for 30–60 min. The cells were disrupted using a Branson Ultrasonics Sonifier 250 on ice, with a very high energy microtip probe (Output control: 5, amplitude: 40%, 5 sec on, 25 sec off, 10 min). The lysate was then centrifuged in a SORVALL LYNX 6000 centrifuge (10,000 × *g*, 45 min), and the supernatant (crude fraction) was collected and used for the enzyme assay. Empty NiCo21 pRARE2 (DE3) cells were processed similarly and the crude fraction was utilized as the control to assess background enzyme activity. SDS-PAGE was used for qualitative analysis and the quantity was accessed Nanodrop at an absorbance ratio of 260/280 (and default protein absorbance values for 0.1%, i.e., 1 mg/mL).

### 2OTS - Tryptamine assays and metabolite profiling

The enzymatic activity of the 2OTS was assessed *in vitro* using the crude enzyme (total vol: 50 µL). The assay comprised the substrate tryptamine (250 µM, dissolved in methanol), FAD (10 µM), NADPH (500 µM), 25 µL assay buffer (100 mM Tris-HCl buffer pH 7.5, 10% glycerol, 10 mM DTT), crude 2OTS enzyme (5 mg), and Milli-Q water. The reaction mixture was thoroughly mixed, and the crude enzyme was added last to start the reaction. As controls, assays without cofactors (NADPH, FAD), and enzyme, were done. Additionally, the background activity of the expression host cell (empty vector) was tested. All the assays were carried out in triplicates. The reaction was incubated at 30 °C (300 rpm), after which the reactions were stopped by adding methanol to a final concentration of 60%. The mixture was further centrifuged to avoid any precipitation in the sample. LC-MS samples were prepared (20% methanol, 0.1% formic acid) and analysed by ultra-high performance liquid chromatography-quadropole time-of-flight mass spectrometry (UHPLC-qTOF-MS) as described below. Data analysis was performed in Bruker Compass DataAnalysis (v. 4.3) software (Bruker Daltonik GmbH 2014, Billerica, Massachusetts, USA).

### Substrate feeding in *Hordeum vulgare*

In this experiment, barley (*H. vulgare)* cultivar RGT Planet was fed with different substrates to investigate a potential metabolic response. The plants were cultivated as described previously. At 1.5 leaf stage (BBCH 11;Lancashire et al., 1991), the first developed leaf was fed with 100 µM solution of either tryptamine, serotonin, or 2-oxo-tryptamine. Control plants were fed with demineralized water. The first forming leaf was infiltrated from the abaxial side of the leaf with a needleless 1 mL syringe. 24 h post feeding, three biological replicates for each substrate were sampled, flash-frozen in liquid N_2_ and stored at −80 °C until metabolite extraction.

### Metabolite extraction and sample preparation

Frozen leaf tissue was homogenized using a TissueLyzer (Qiagen) and pre-cooled inserts (30 s, 30 Hz). Metabolites were subsequently extracted from the homogenized plant tissue using a cold extraction buffer (85% methanol, 0.5% formic acid, 250 µM caffeine (Sigma) and 250 µM sambunigrin [synthesized in-house], as internal standards). Immediately after adding the extraction buffer, samples were vortexed and then boiled (5 min at 80 °C). Samples were cooled on ice (10-20 min) and centrifuged (5 min, 4 °C and 20.000 x *g*). Prior to UHPLC-qTOF-MS, the supernatant was diluted five times and filtered through a 0.22 µM Millipore GV filter (Merck Millipore, Burlington, Massachusetts, USA) by centrifugation (5 min, RT and 3000 rpm) and transferred to glass vials with 0.1 mL inserts (Mikrolab, Aarhus). Filtered extracts were stored at 4 °C until further analysis (< 24 hours). Throughout the sample preparation, samples were randomized to minimize bias.

### Metabolomic analysis using Ultra-High Performance Liquid Chromatography – Mass Spectrometry

#### Ultra-High Performance Liquid Chromatography

Filtered methanol extracts were separated on a Dionex Ultimate 3000RS ultra-high performance liquid chromatography (UHPLC; Thermo Fisher Scientific, Waltham, Massachusetts, USA) system with a diode array detector. The autosampler and column were maintained at 10 °C and 40 °C, respectively. Four µl of extracts were injected onto and separated using a Phenomenex Kinetex® column (100 Å, 1.7 µm C18, 150 x 2.1 mm, Phenomenex Inc., Torrance, California, USA). Leucine-Enkephalin ([M+H]^+^ = 556.2771) was used as reference calibrant and injected in the beginning and end of the sample sequence to monitor sensitivity and performance of the UHPLC system. A binary solvent system consisting of water (A) and acetonitrile (B) both supplied with 0.05% (v/v) formic acid was used for the gradient elution at a flow rate of 0.3 mL per min. The gradient elution profile was as follows: 0.0-2.0 min, 2% B; 2.0-15.0 min, 2-40% B; 15.0-17.0, 40-100% B; 17.0-18.0, 100% B; 18.0-20.0, 100-2% B; 20.0-25.0, 2% B. The total run time for each sample was 25 min. In between each injection, the analytical column was allowed to re-equilibrate for 5 min.

The system stability was assessed using quality control (QC) samples consisting of pooled aliquots of all plant extracts. The QC samples were run in triplicate in the beginning and end of the total run as well as recurrently throughout acquisition. Injection order of extracts was randomized in order to minimize measurement bias. Samples were analyzed in batches of ten with each batch being separated by a system blank (100% methanol) and a QC sample. Additionally, a process blank (PB) sample was run in the beginning and end of the sample providing information on possible contaminants related to sample preparation. The PB sample was prepared following the same protocol as described but without any biological material.

#### Mass spectrometry

The UHPLC system was connected to a compact^TM^ quadropole time-of-flight mass spectrometer (q-TOF MS) (Bruker Daltonics, Billerica, Massachusetts, USA). Electrospray ionization was used for ionization of the analytes in positive mode with an ion spray voltage maintained at 4000 V. End plate offset was set to 500 V. Nitrogen was used as the desolvation, nebulizing gas, and the collision gas. The desolvation gas temperature and flow rate was set to 250 °C and 8 L/min, respectively. The nebulizing gas pressure was 2.0 bar. Mass (both MS1 and MS2) spectra were acquired in a scan range of 50 to 1400 *m/z*, scan time 0.3 s and collision energy at 7.0 eV. Active exclusion number was set to 3, and the precursor reconsideration time at 0.2 min; thus, after 3 MS/MS spectra had been recorded for a specific precursor ion it was excluded for analysis for the set time. The sampling rate was 4 Hz. Sodium formate (HCOONa) was used for internal mass calibration.

### Matrix-Assisted Laser Desorption Ionization-Time of Flight Mass Spectrometry (MALDI-TOF MS)

#### Embedding, sectioning, and freeze-drying

Fresh barley leaves were embedded using aqueous mixture of hydroxypropyl-methylcellulose and polyvinylpirrolidone (Mw 360kDa) (7.5 % (w/v) + 2.5 % (w/v), respectively) and sectioned in cross-section orientation in a Leica CM3050S Cryostat at –25 °C using Kawamoto’s film method (Kawamoto & Kawamoto, 2021) adapted for plant samples (Montini et al., 2020) to prepare 14 µm thick sections. Transparent adhesive cryo-tape for frozen surfaces (Labtag, Laval, Canada) was used to obtain intact sections which were then mounted onto glass slides using double-sided FrozenSTUCK cryo-tape (Labtag, Laval, Canada). The sections were freeze-dried under vacuum overnight and stored under vacuum, at room temperature in black boxes.

#### Matrix deposition

MALDI matrix 2,5-dihydroxybenzoic acid (DHB) was applied as 7.5 mg/mL DHB solution in ethanol/water/formic acid (90:10:0.1) by spraying using HTX TM-Sprayer (HTX Technologies, Chapel Hill, North Carolina, USA). The spraying parameters were the following: flow rate 0.1 mL/min, nozzle temperature 60 °C, nozzle velocity 1250 mm/min, nozzle height 40 mm, line spacing 2 mm, number of passes 20, done in a horizontal movement pattern. Gas flow was set to 3 L/min with gas pressure kept at 10 psi. Sections with applied matrix were analyzed on the same day.

#### Mass Spectrometry Imaging

Section with deposited matrix were imaged using timsToF-fleX MALDI-2 instrument (Bruker Daltonics, Billerica, Massachusetts, USA) with post-ionization applied (MALDI-2). The imaging was performed with 20 µm spatial resolution, with a scan range of m/z 50-1000, using 30 shots/pixel, “Single” laser geometry with active bean scan applied and trigger delay of 10 µs for the MALDI-2 postionization. Selection of the analyzed area on the sample was done using Bruker Compass flexImaging 7.6 (Bruker Daltonics, Billerica, Massachusetts, USA). Images were generated using SciLS Lab version 2024a Pro software (Bruker Daltonics, Billerica, Massachusetts, USA) with ±0,005 Da error interval to the exact mass. Data were normalized using Root Mean Square (RMS) normalization.

### Structural elucidation of 2OT

#### Bulk metabolite extraction of infected barley leaf tissue

Leaf material from barley *cv.* RGT Planet inoculated with *P. teres* was sampled in bulk 7 dpi and snap-frozen in liquid N_2_. Subsequently, frozen leaf tissue was homogenized using pre-cooled pestle and mortar. Metabolites were extracted using a hot extraction buffer (70% methanol, 65 °C). The buffer was added in a 10:1 (w:w) ratio to the homogenized sample and mixed for 2 min. The extraction was filtered through a Büchner funnel coupled with a filter (Teknisk papirfilter nr 118, Ø 85 mm, MIKROLAB – FRISENETTE, Denmark) and subsequently centrifuged (5 min, 4200 rpm and 7 °C) in a tabletop centrifuge (5424 centrifuge, Eppendorf). Subsequently, the supernatant was transferred to a glass bottle and concentrated using a rotary evaporator (heated at 35 °C, 150 mbar, cooled at −6 °C). Hereafter, the aqueous sludge was transferred to a new glass vial and freeze dried. The complete pellet was dissolved in 10% (v:v) ethanol to match the starting conditions during Flash Chromatography purification.

#### Isolation and purification

2OT was isolated using a PuriFlash 5.250 coupled with an ELSD detector using a Uptisphere® Strategy^TM^ PF-15C18HQ-F0025 Interchim® column (100 Å, 15 μm C18, 224 x 36 mm; Interchim, Montluçon, France). Fractions were collected in 20 mL glass tubes. A binary solvent system consisting of water (A) and ethanol (B) was used for the gradient elution at a flow of 15 mL per min. The gradient elution profile was as follows: 0.0-15.0, 5-40% B; 15.0-17.0, 40-100% B; 17.0-25.0, 100% B; 25.0-28.0, 100-5% B; 28.0-32.0, 5% B. During elution, fractions were collected at three different wavelengths: 280 nm, 240 nm, and 210 nm with 0.3 mAU, 10 mAU, and 10 mAU threshold, respectively. In addition, manual collection was done to make sure to precisely isolate the fractions of interest in different glass tubes. Collected fractions were combined and ethanol was evaporated under nitrogen flow. The remaining aqueous sample was freeze-dried and subsequently redissolved in 80% methanol and analyzed via UHPLC-qTOF-MS/MS. Structure elucidation was performed by NMR spectroscopy.

#### Nuclear Magnetic Resonance

NMR samples were dissolved in DMSO-d6 or CDCl3 (calibrated to δ_H_ 2.50 ppm and δ_C_ 39.50 ppm or δ_H_ 7.26 ppm and δ_C_ 77.16 ppm, respectively) and 1D ^1^H, ^13^C and DEPT-135 spectra for synthetic compound references were acquired at 300 K on a Bruker Avance III spectrometer (^1^H operating frequency of 599.58 MHz) equipped with a Bruker SampleCase autosampler and a cryogenically cooled ^13^C/^1^H DCH 5-mm probe-head (Bruker Biospin, Karlsruhe, Germany) using standard pulse sequences and parameters (16 scans and a relaxation delay of 1 s for ^1^H and 256 scans and 2 s for ^13^C). 1D ^1^H, HSQC and HMBC spectra for minor metabolites (and 2OT reference) were acquired at 300 K on a Bruker Avance III spectrometer (^1^H operating frequency of 600.13 MHz) equipped with a Bruker SampleJet sample changer and a cryogenically cooled gradient inverse triple-resonance 1.7 mm TCI probe-head (Bruker Biospin, Karlsruhe, Germany) using standard 1D and 2D experiments (64 scans and 1730×256 data points for HSQC and 128 scans and 2048×256 data points for HMBC). Data was processed using Topspin ver. 4.4.0 (Bruker).

### Data processing

The acquired raw spectral data was calibrated using the Bruker Compass DataAnalysis (v. 4.3) software (Bruker Daltonik GmbH 2014, Billerica, Massachusetts, USA) and subsequently converted to open-source format *.mzML* files using msConvert, which is a part of the ProteoWizard Toolkit (Chambers, Maclean et al. 2012, Adusumilli and Mallick 2017). Hereafter, precursor *m/z* values were corrected by applying a freely available script on the *.mzML* files (Breaud, Lallemand et al. 2022). The script was accessed on https://github.com/elnurgar/mzxml-precursor-corrector 25th of March 2025. A PCA was performed, which showed narrow clustering of QC pooled samples and blanks, indicating overall reliable spectral data acquisition throughout the analysis (Figure S1).

#### MZmine

The converted files were further processed using MZmine4 (v. 4.0.3) (Pluskal, Castillo et al. 2010, Schmid, Heuckeroth et al. 2023). A signal intensity noise cutoff of 200 and 100 was applied to MS1 and MS2 data, respectively. For scans in retention time window 0.5-15.0 min, chromatograms were built using the ADAP chromatogram algorithm with an *m/z* tolerance of 0.005 *m/z* (20 ppm) and minimum absolute height of 2000. Post feature detection, smoothing was applied using the Savitzky-Golay algorithm (Savitzky and Golay 1964). The Local minimum feature resolver algorithm was used to deconvolute the generated extracted ion chromatograms. Isotopes were detected using the 13C isotope filter whereafter the isotopic peaks finder was applied. The Join aligner algorithm was used to align the feature list with an *m/z* and retention time tolerance of 0.006 *m/z* (10 ppm) and 0.1 min, respectively. To account for missing values in the aligned feature table due to data preprocessing, gap-filling was applied using the Peak finder algorithm followed by the Duplicate feature filter (New average mode) to avoid misaligned feature list rows post gap-filling. The resulting aligned feature list were subjected to both Spectral/Molecular and Ion Identity networking and exported using the GNPS-FBMN/IIMN and SIRIUS/CSI-FingerID modules (Wang, Carver et al. 2016, Nothias, Petras et al. 2020, Schmid, Petras et al. 2021). A detailed description of all parameters used for MZmine4 processing in this study can be found in Table S5.

#### Feature-Based Molecular Networking

The GNPS export files (*.csv* quant table, *.mgf* MS/MS spectral summary, and .csv additional edges) generated in MZmine4 were uploaded to the GNPS2 platform. A feature-based molecular network (FBMN) was generated for the entire spectral dataset using the FBMN workflow (v. 2024.10.09) (Nothias, Petras et al. 2020). The precursor ion tolerance was set to 0.02 Da with a fragment ion tolerance of 0.02 Da. For spectral similarity, the cosine score threshold was 0.7 with at least 6 matched fragment ions. For the spectral library search, the minimum cosine similarity score required was 0.7 with a minimum of 6 matching peaks. When computing FBMN for the entire spectral dataset, a task was submitted both with and without applying search for analogs to library spectra in data (Top-K set to 1). A description of all parameters used for FBMN in GNPS2 can be found in Table S5. Links for the FBMN jobs will be made public once the manuscript is published; until then, they can be shared upon request. The generated FBMN was visualized in the Cytoscape network visualization software (v. 3.10.2) (Shannon, Markiel et al. 2003).

#### SIRIUS

The SIRIUS *.mgf* file containing spectral information for each feature generated in and exported from MZmine4 was processed with SIRIUS (v. 5.8.6) (Dührkop, Fleischauer et al. 2019). The instrument profile was set to q-TOF and the MS2 mass accuracy to 10 ppm. Remaining parameters were left as default. The prediction of the molecular formulas was improved by applying ZODIAC with default parameters (Nothias et al., 2020). Structure predictions was done using CSI:FingerID and the COSMIC workflow (*Fallback Adducts*: [M + H]+, [M + K]+, [M + Na]+, [M + H3N + H]+, [M + CH4O + H]+, [M + C2H3N + H]; *Search DBs*: Bio Database and Natural Products) (Dührkop et al., 2015; Hoffmann et al., 2022). CANOPUS was applied for *de novo* chemical compound class predictions using the NPClassifier taxonomy (Dührkop et al., 2015; Kim et al., 2021). Compounds with a mass above 850 Da were excluded from analysis due to computational limits. A description of the parameters used for SIRIUS can be found in Table S5.

#### MS-Dial

LC-MS data from methanol extracts of barley leaves exogenously treated with tryptamine, 2OT, serotonin, or water (control) were processed using MS-DIAL version 5.1 (Takeda et al., 2024; Tsugawa et al., 2015). Key parameters included MS1 and MS2 tolerances of 0.01 Da and 0.025 Da, respectively. Peak detection was performed with a minimum peak height of 3000 amplitude and a mass slice width of 0.1 Da. MS2Dec was configured with a sigma window value of 0.5 and an MS/MS abundance cut-off of 0. For feature alignment, retention time tolerance was set to 0.1 min and MS1 tolerance to 0.025 Da. Features showing a fold change (Max/Min) >2 and a p-value <0.05 relative to the control were manually inspected, annotated when possible, and cross-referenced with the FBMN derived from pathogen-infected barley leaves.

### Statistical analyses

#### Blank subtraction and imputation

Features derived from blank samples were removed from the MZmine4 generated feature table using the Jupyter Notebook (R script) provided by (Pakkir Shah et al., 2023). The notebook was executed in the Google Colab environment. Blank subtraction was done with a cutoff filter of 0.3, i.e. if the ratio of average intensity for any feature in blanks to samples were above 0.3, the feature was removed. Zeros present in the blank-removed feature table were randomly replaced by created random values between 0 and the minimum value in the blank-removed feature table. In R (v 4.4.3), feature intensities were normalized to mg fresh weight in the samples.

#### Random Forest analysis

The supervised machine learning algorithm Random Forest (RF) (Breiman, 2001) was used to evaluate whether the metabolic space detected in our dataset could be separated by the treatments (control versus infected) at 4 dpi. The precision of the separation by RF is evaluated with the out-of-bag (OOB) prediction error. Furthermore, RF was used to obtain a list of variables most important for separation between treatments. The analysis was done in R (v 4.4.3) using the ‘randomForest’ function from the *randomForest* package (Liaw & Wiener, 2002) with *n*_tree_ = 10.000 and *m*_try_ = 30. The results were visualized using the *ggplot2* R package (Wickham, 2016).

#### Principal component analysis

The unsupervised learning method principal component analysis (PCA) was performed on the blank-subtracted, imputed, and normalized metabolomic dataset. The analysis was done in R (v 4.4.3) using the ‘pca’ function from the *FactoMineR* package (Francois Husson et al., 2006) with *ncomp* = 5, *scale* = TRUE, and *center* = TRUE. The results were visualized using the R packages *ggplot2* (Wickham, 2016) and *mdatools* (Kucheryavskiy, 2014).

### Metabolite annotation

An in-house spectral library consisting of 11 reference compounds (commercially sourced or in-house synthesized) analyzed with the same UHPLC-qTOF-MS/MS method as described above was used for metabolite identification. The reference compounds used within this study are listed in Table S6. The in-house spectral library was generated in MZmine4 (v.4.0.3) using the Local compound database search and Batch spectral library generation modules.

All compounds annotated in this study are listed in Table S1. Information on level of confidence for the identification of the individual metabolites is included. The levels of identification referred to follows the guidelines by Metabolomics Standards Initiatives (Sumner et al., 2007) and the proposed modifications by (Schymanski et al., 2014); (1) structure confirmed by m/z, retention time and MS/MS matching compared to a reference standard, (2) putative structure identified based on MS/MS matching to library or literature spectrum, (3) tentative structure candidate due to insufficient information to determine one exact structure (i.e. positional isomers or top hit from prediction in the *in silico* tool SIRIUS), (4) molecular formula, and (5) exact mass only.

### Chemical synthesis of *N*-hydroxytryptamine and its analogue 3-(2-aminoethyl)-1H-indol-1-ol

*N*-hydroxytryptamine was chemically synthesized according to a published procedure (Hee et al., 1992). 3-(2-Aminoethyl)-1H-indol-1-ol was chemically synthesized from the commercially available tryptamine, thus the primary amino group was first protected, then NH of the indole moiety was hydroxylated using m-chloroperbenzoic acid followed by deprotection of the primary amino group to provide the target molecule (unpublished data). The identities of the synthesized compounds were ascertained by NMR spectroscopy and agreed with reported data.

### Quantitative determination of plant metabolites using standard curves

The concentration of tryptophan, tryptamine, serotonin, hydroxylated tryptamine derivative [2], and 2OT in the plant extracts was determined using a prepared standard curve. Standard solutions of authentic tryptamine (Sigma-Aldrich), serotonin (Sigma-Aldrich), and 2OT (BLDpharm) were prepared in methanol at concentrations ranging from 0.1 to 50 µM. The standard solutions were analyzed using the same UHPLC-MS/MS method as described previously. A standard curve was constructed by plotting concentration against peak area, and the linear regression was employed for quantification. The standard curve for serotonin was used to quantify abundance of tryptophan and hydroxylated tryptamine derivative [2] since the ionization efficiency in the system was the same. Reported values are means of the biological quintuplicates, which were extracted and analyzed for each sample type.

### RNA extraction and sequencing

Total RNA was extracted from 25 to 60 mg leaf tissue using the Spectrum^TM^ Plant Total RNA kit (Sigma-Aldrich, US), following the manufacturer’s protocol A. The optional on-column DNase digestion was done using The On-Column DNase I Digestion set (Sigma-Aldrich, US). Transcriptomes were prepared from 36 samples consisting of four biological replicates per treatment (infected and mock-control) and timepoint (1, 2, 4, and 7 dpi) representing nine distinct sample conditions, including the T0 no-treatment control. RNA quality and integrity were assessed by NanoDrop and Bioanalyzer 2100/GX. Library preparation and sequencing were performed by Biomarker Technologies (BMK) GmbH using an Illumina NovaSeq X (Illumina, San Diego, CA) to generate 30M paired-end reads of 150 bp (9 Gbp per sample). Raw data files will be made public once the manuscript is published; until then, they can be shared upon request.

The raw sequencing data was processed by Sequentia Biotech®. The FastP tool was used for read quality filtering and trimming ensuring that low-quality bases and adapter sequences were removed. Subsequently, the software Kraken was used to classify the trimmed reads in order to remove unwanted rRNA-specific sequences. The high-quality reads were then aligned against the barley (*H. vulgare*) *cv.* RGT Planet genome (v2) (Jayakodi et al., 2024) using STAR with default parameters. Gene expression was quantified using FeatureCounts and normalized with Fragments Per Kilobase Million, Trimmed Mean of M values, and TPM normalization. Lowly expressed genes were removed in R using the HTSFilter package leaving 20065 filtered genes. These were used as input for differential gene expression analysis, which was done using the DeSeq2 R package. Genes with a p-value < 0.05 and p.adj. < 0.05 were considered statistically significant. The differential expression analysis was done comparing the two different treatments (infected versus control) at all individual time points (T1, T2, T4, and T7) as well as through the course of all time points.

### Bioactivity assay

Potential bioactivity of serotonin and 2OT was tested against the fungal phytopathogen *P. teres*. Hordenine was included as positive control, and demineralized water (2% methanol, 0.2% DMSO) as negative control. The different compound and concentration combinations were tested in triplicate. *P. teres* was prepared as described under “Pathogen Inoculum Preparation”. Spore concentration was determined using a haemocytometer. The bioactivity of the compounds of interest was tested at 1 and 5 mM. The final reaction had a spore concentration of 3000 spores/mL, 2% methanol, and 0.2% DMSO. Reaction mixture was vortexed, and subsequently incubated (RT, 200 rpm) for 24 h. Hereafter, 200 µL were plated on V8 agar plates using sterile glass beads. After five days at RT, fungal growth was visually inspected.

### Data records/availability

The raw *.d* and converted *.mzML* UHPLC-qTOF-MS/MS data files were uploaded to the MassIVE repository under the accession number MSV000097403 in designated folders “Raw” and “mzML”, respectively. This repository also contains (1) a *.txt* metadata file with information related to each sample on treatment, time point, fresh weight sample mass [mg], and cultivar, (2) ReDU metadata (Jarmusch et al., 2020) enabling co- and re-analysis with publicly available data in the GNPS environment, (3) a *.csv* file containing combined annotation results from GNPS, SIRIUS, CANOPUS, and MZmine, (4) a document with information on MZmine4, FBMN, and SIRIUS data processing parameters, and (5) the *.cys* Cytoscape file for the visualization of the FBMN. The MassIVE accession will be made publicly available once the manuscript is published; until then, access can be shared upon request. The same applies to the GNPS2 hyperlinks for the FBMNs generated within this study as well as the RNA-seq raw data files.

## Acknowledgements

Bachelor students Kristian Siig Hessel, Karen Fjordside Miltersen, Philip Kierstein Kjærside and Ida Agerbirk Rytter for initial characterization of barley CYPs. Mads Eggert Nielsen for providing the *B. hordei* infected leaf tissue and Kenneth Madriz Ordenana for providing the *P. syringae*. We thank David R Nelson for naming the cytochromes P450 in this study.

## Funding

This work was supported by the Independent Research Fund Denmark (Grants 1051-00083B & 1131-0002B) and the Novo Nordisk Foundation (Grant No. 0054890) awarded to EHJN. MV acknowledges the Marie Skłodowska-Curie Individual Fellowship (MSCA grant agreement No. 101110417).

## Author contributions

EHJN and ST conceived the project. JMC and EHJN designed the project, which was supervised by EHJN and MB. HLJ, MMTA and JMC performed the infection studies. MMTA and JMC performed the metabolomic and transcriptomic analyses. MP and JG performed the 2OT purification. MSM performed the chemical synthesis while LK and DS performed the NMR. JMC, MMTA and EHJN wrote the paper with comments from all authors.

## Notes

### Competing Interest Statement

The authors have declared no competing interest.

