## Supplementary Material for "FMO and CYP monooxygenase families determine the metabolic flux of hydroxylated tryptamine derivatives in barley (*Hordum vulgare*) following pathogen infection"

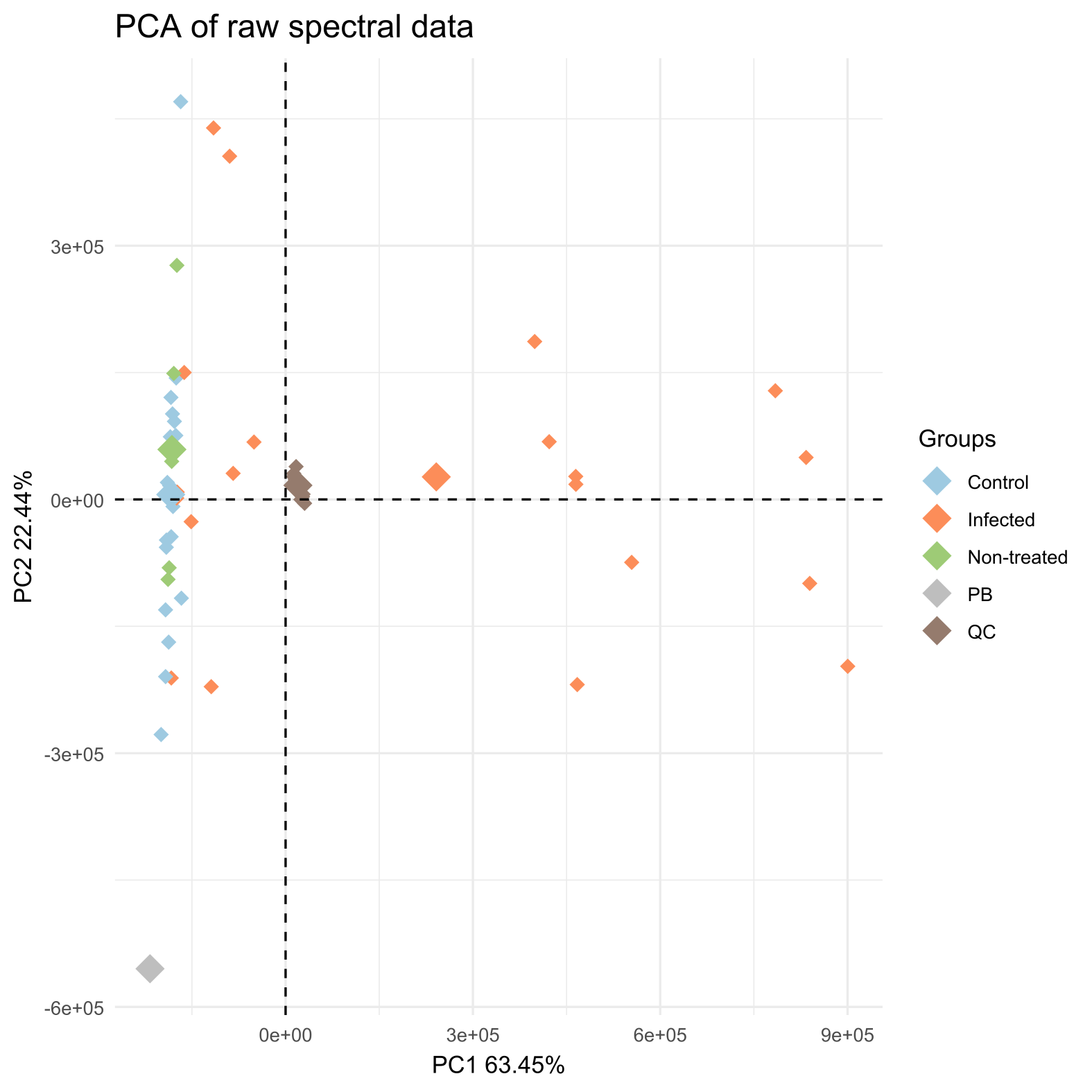

**Figure S1: Assessment of reliability and quality of UHPLC-qTOF-MS/MS data acquisition.** A principal component analysis (PCA) of the acquired full spectral dataset showed narrow clustering of quality control (QC) pooled samples (brown) and blanks (PB; grey).

**Figure S2: Relative abundance of detected features in the chemical space of infected and non-infected barley leaves.** Global feature-based molecular network (FBMN) visualizing the detected metabolic landscape of barley *cv.* RGT Planet. The 38 largest molecular families (i.e. ≥ 3 nodes) in the network are numbered. Pie-charts indicate relative abundance of individual features in control (blue) and *P. teres* infected (orange) leaf samples across entire spectral dataset.

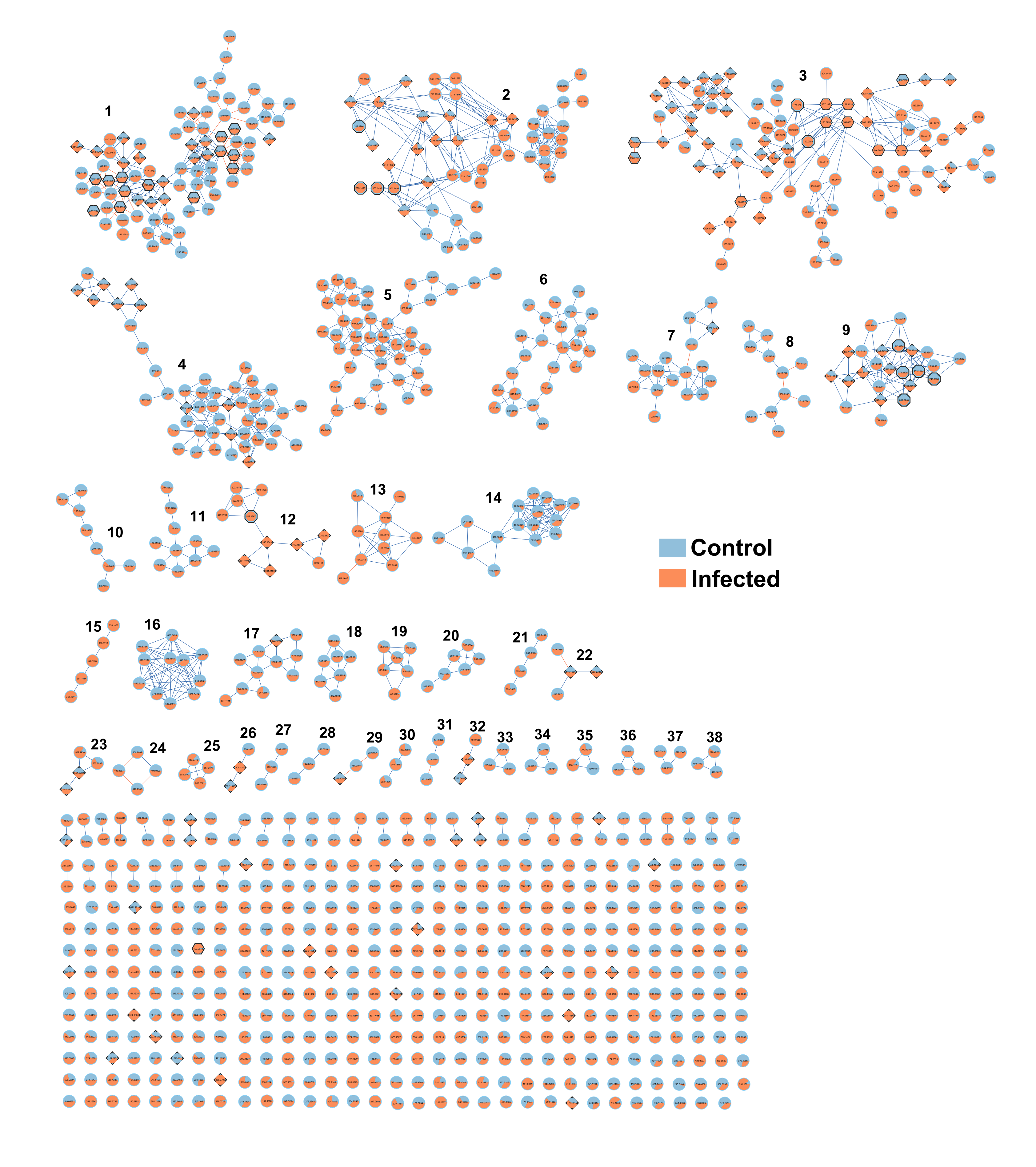

**Figure S3: Principal component analysis of metabolites extracted from *P. teres* infected, mock control, or non-treated barley leaves across experimental time course.** The principal component (PCA) scores plot for PC1 and PC2 **(A)** show strong separation of infected (orange) from mock control (blue) and non-treated (green) samples from 4 days post inoculation (dpi) with *P. teres*, while the PCA scores plot for PC3 and PC4 **(B)** show a distribution of the different sample types driven by ontogenetic development. The PCA loadings plots **(C) and (D)** show the features driving the distribution of the samples.

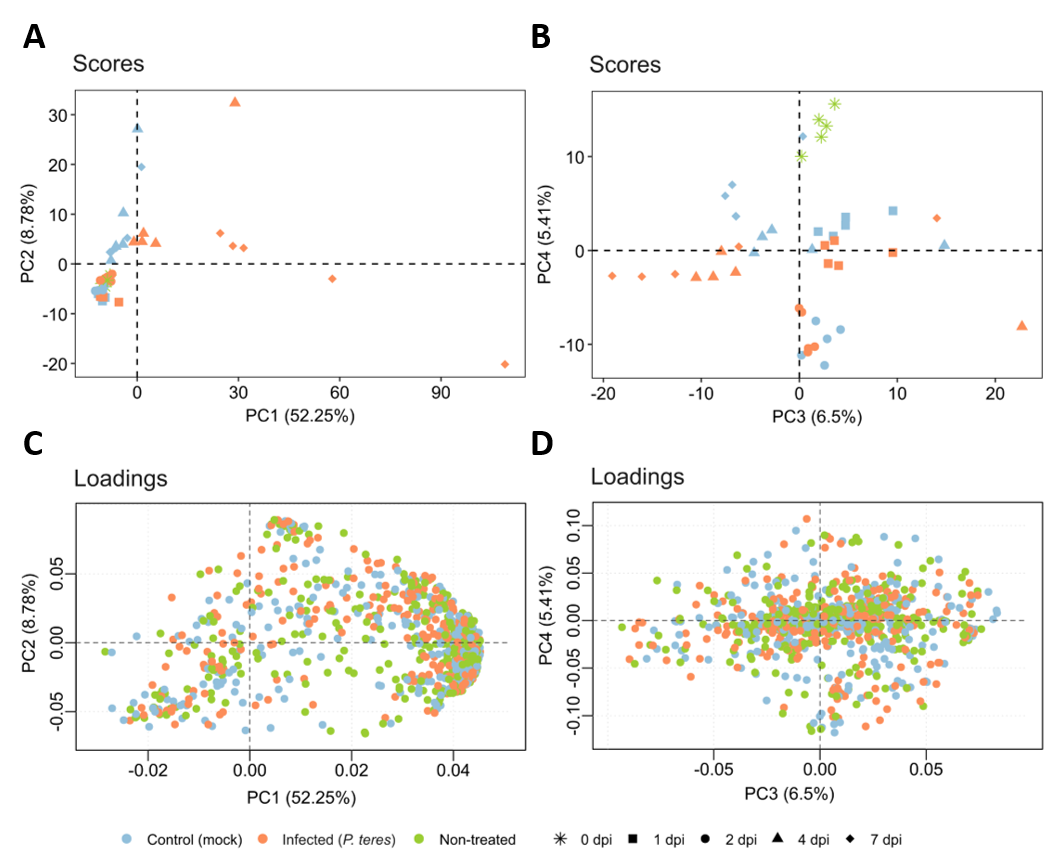

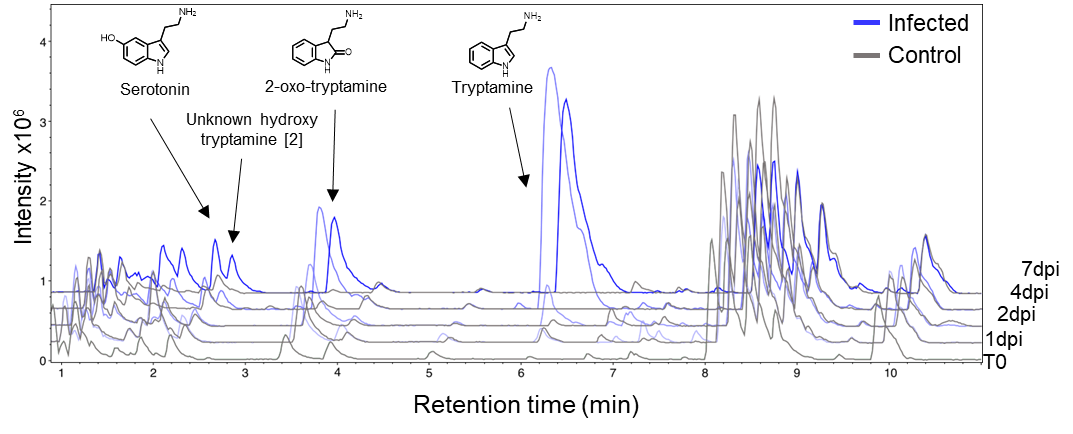
**Figure S4: Chromatographic comparison between infected and control through experimental time course.** Base peak chromatograms of representative methanol extracts of barley leaves infected with *P. teres* (blue) and control (treated with water; gray) at 1, 2, 4 and 7 days post inoculation (dpi). Serotonin, 2OT and tryptamine are labelled and verified by authentic standards.

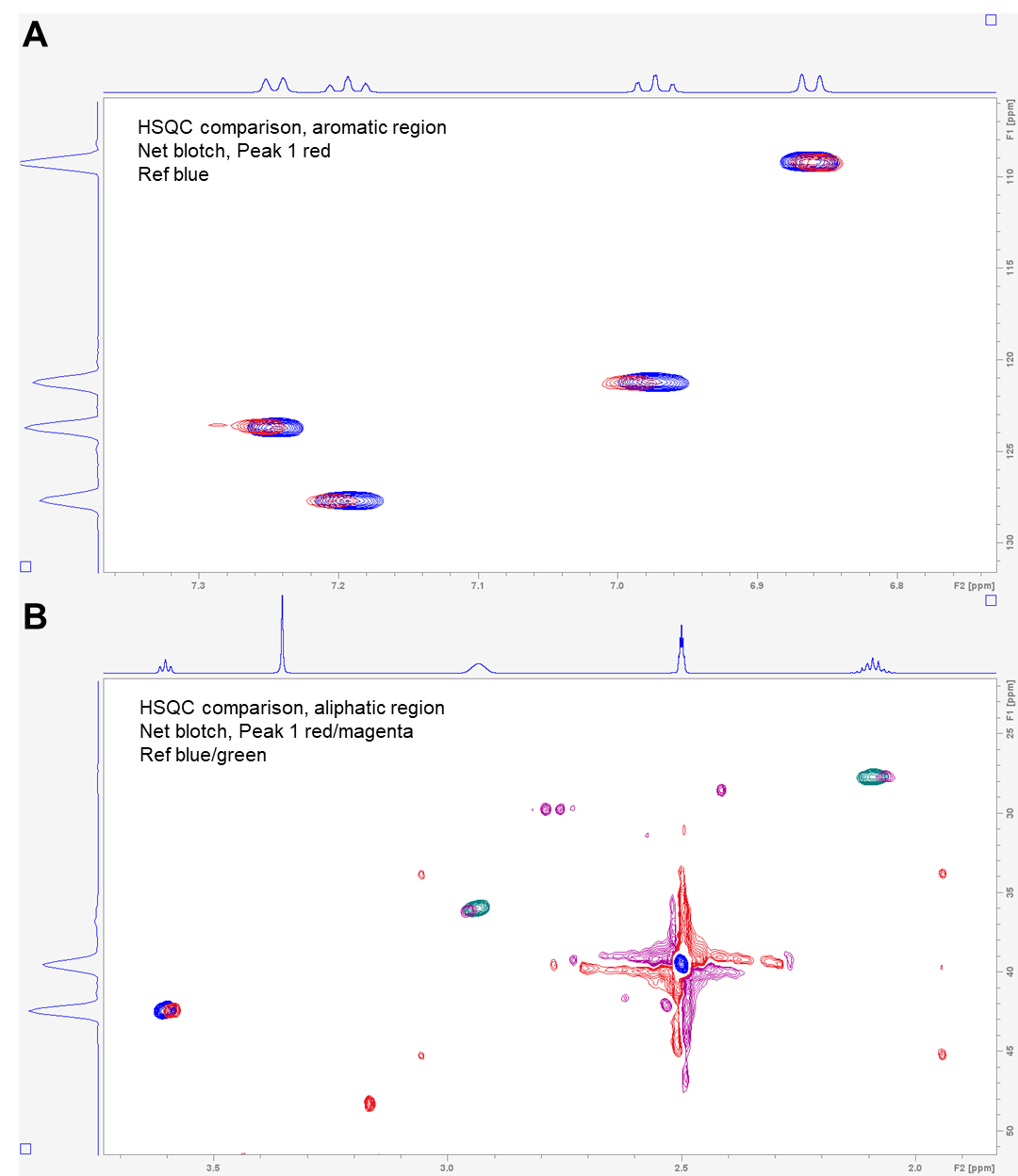

**Figure S5:** **HSQC NMR fingerprint comparison** **of 2OT authentic standard and purified 2OT from *P. teres* infected barley leaf sample.** This figure shows the HSQC NMR spectra of the 2OT authentic standard reference (blue and green) and 2OT purified from a *P. teres* infected barley leaf sample (net blotch; red and magenta) for **(A)** the aromatic, and **(B)** the aliphatic region of the compound. The correlations in the HSQC spectra confirms that the purified compound from the infected barley leaf sample matches the authentic standard.

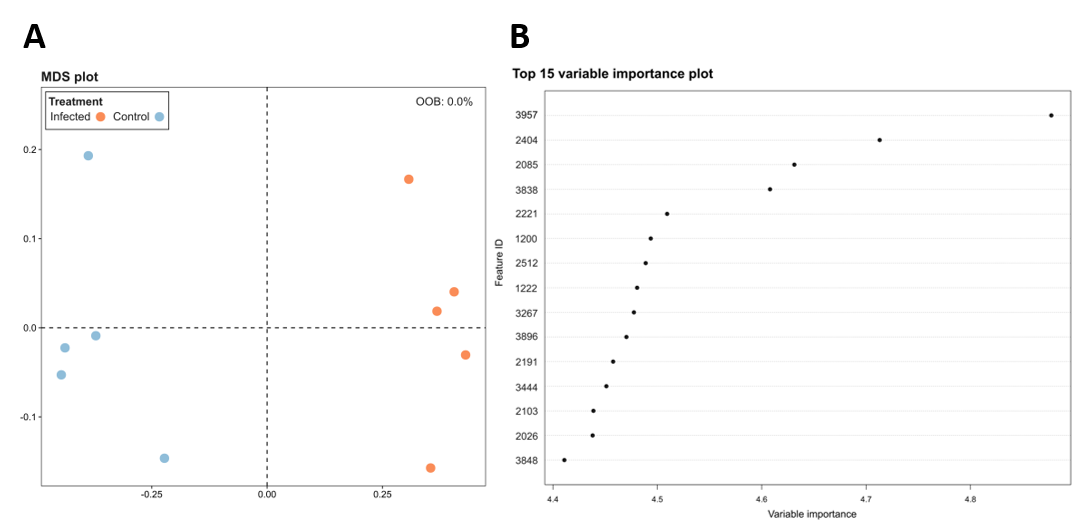

**Figure S6: Random Forest analysis of barley *cv.* RGT Planet infected with *P. teres* at 4 dpi. (A)** Metric multi-dimensional scaling (MDS) plot of the proximity matrix obtained from Random Forest (RF) analysis showing significant differences in the metabolome of control (blue) and infected (orange) barley leaves at 4 dpi with an out-of-bag (OOB) prediction error of 0%. **(B)** Top 15 important variables explaining the differences in the metabolome obtained from RF analysis are shown, and information on these variables/features are provided in Table S2.

**Table S2: Top 15 important variables explaining metabolomic differences in infected versus mock-control barley leaves at 4 dpi by random forest analysis.** Additional information on the top 15 variables/features contributing to the observed metabolic differences between *P. teres* infected and mock-control barley leaves at 4 dpi obtained from Random Forest (RF) analysis (Figure S6). Abbreviations: RT, retention time; MF, molecular family in feature-based molecular network; NPC, NPClassifier taxonomy (Kim et al., 2021); VIP, variable importance.

| **Feature ID** | ***m/z*** | **RT** | **MF** | **Putative annotation** | **Annotation level** | **NPC Pathway**  **(CANOPUS)** | **VIP** |
| --- | --- | --- | --- | --- | --- | --- | --- |
| 3957 | 477.176 | 14.913 | 12 | NA | NA | Shikimates and Phenylpropanoids | 4.877 |
| 2404 | 305.16 | 7.898 | 2 | NA | NA | Alkaloids | 4.713 |
| 2085 | 307.141 | 6.285 | 2 | NA | NA | Shikimates and Phenylpropanoids | 4.631 |
| 3838 | 359.15 | 13.622 | 12 | 3,4,2',4',6'-Pentamethoxychalcone | 2 | Shikimates and Phenylpropanoids | 4.608 |
| 2221 | 202.099 | 7.264 | Doublet | NA | NA | Alkaloids | 4.509 |
| 1200 | 376.137 | 1.457 | Singleton | NA | NA | Shikimates and Phenylpropanoids | 4.494 |
| 2512 | 351.156 | 8.273 | Singleton | NA | NA | NA | 4.489 |
| 1222 | 167.581 | 1.516 | Singleton | NA | NA | NA | 4.481 |
| 3267 | 667.297 | 11.315 | 5 | NA | NA | Amino acids and Peptides | 4.478 |
| 3896 | 353.149 | 14.288 | 2 | N-feruloylserotonin | 3 | Amino acids and Peptides | 4.470 |
| 2191 | 132.045 | 7.111 | Singleton | NA | NA | Alkaloids | 4.458 |
| 3444 | 318.139 | 11.922 | Singleton | NA | NA | Carbohydrates | 4.451 |
| 2103 | 371.03 | 6.415 | Singleton | NA | NA | Carbohydrates | 4.439 |
| 2026 | 161.108 | 6.065 | 3 | Tryptamine | 1 | Alkaloids | 4.438 |
| 3848 | 511.232 | 13.761 | Singleton | NA | NA | Alkaloids | 4.411 |

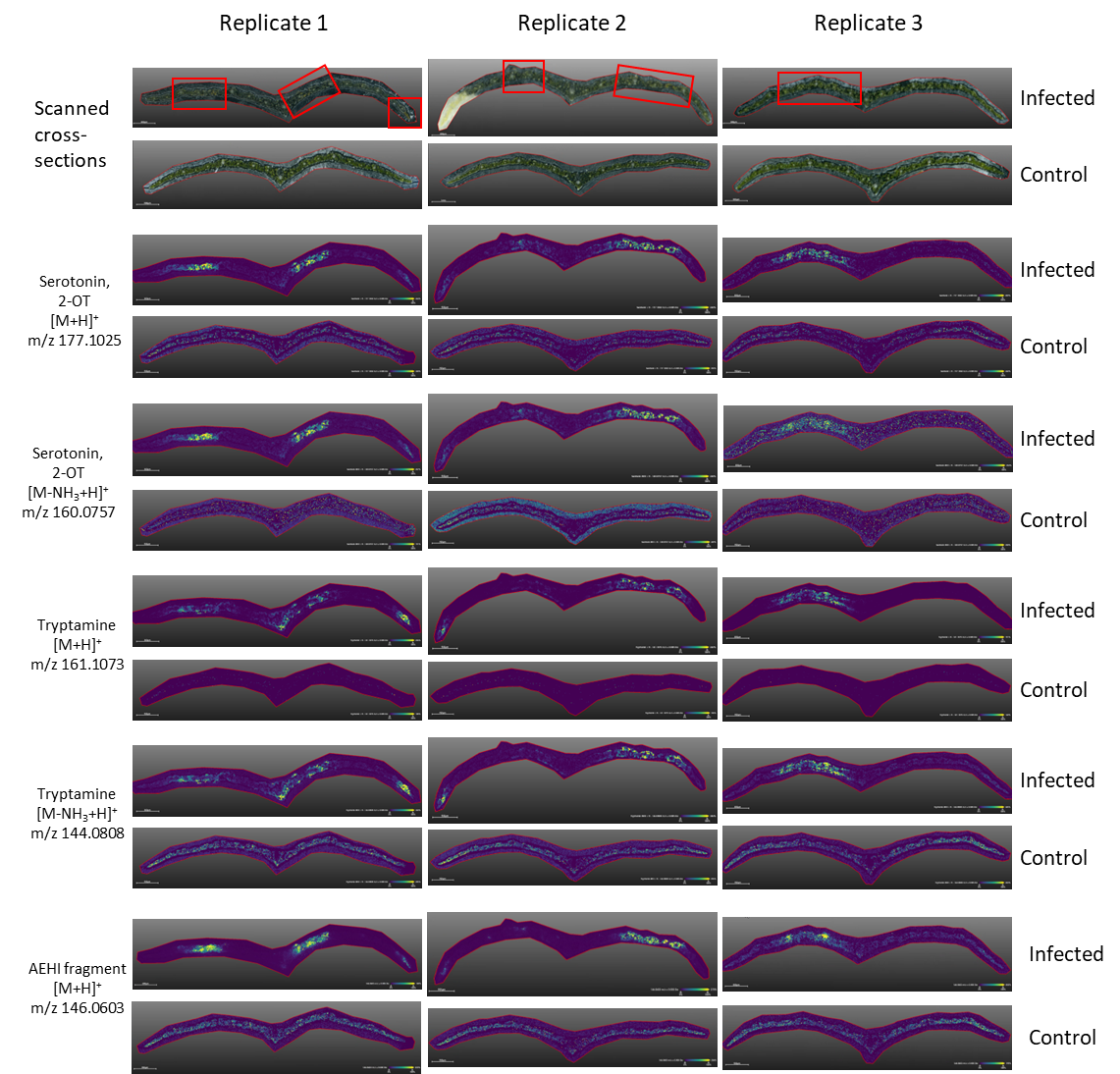

**Figure S7: Localization of tryptamine and hydroxylated tryptamine derivatives (serotonin and 2-oxo-tryptamine (2-OT)) in infected and control samples of barley leaves.** Serotonin and 2OT are detected as isobaric signals corresponding to [M+H]^+^ (*m/z* 177.1025) and [M-NH_3_+H]^+^ (*m/z* 160.0757) ions. Tryptamine signals correspond to [M+H]^+^ (*m/z* 161.1073) and [M-NH_3_+H]^+^ (144.0808) ions. AEHI signals correspond to [M+H]+ (m/z 146.0603). All signals are predominantly detected in infected regions, indicated by red squares. Scanned images of infected and control samples cross-sections are shown in top row of the figure. Color bar shows intensity of detected ions with yellow and blue colors corresponding to higher and lower intensity, respectively. All images are generated with ± 5mDa error interval.

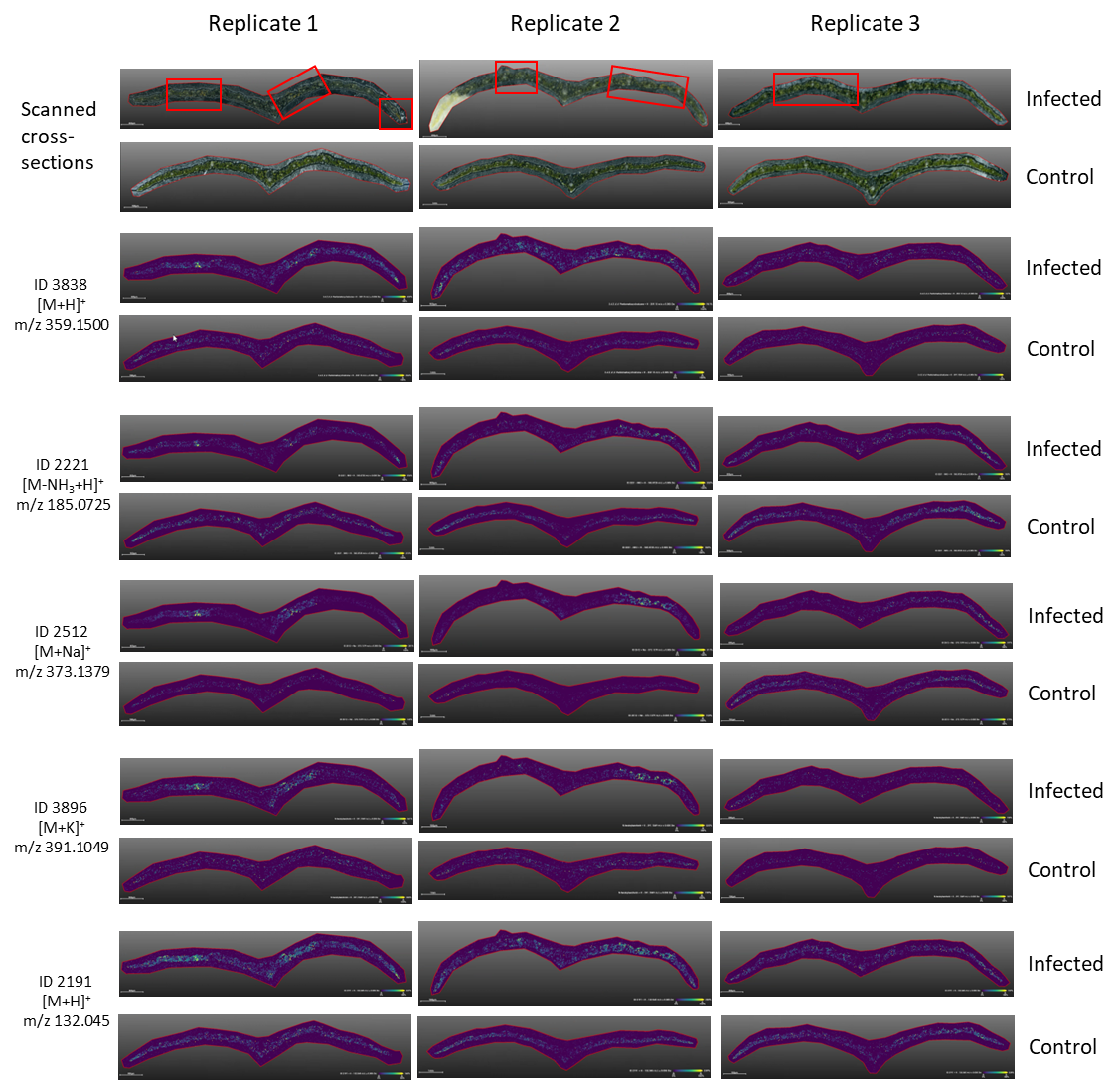

**Figure S8: Localization of phytoalexins identified in Random Forest analysis in infected and control samples of barley leaves.** Each compound is assigned with a unique ID representing the feature ID in the molecular network. The features investigated were identified among top 15 most important variables contributing to the variance between *P. teres* infected and control barley leaves at 4 dpi in a Random Forest (RF) analysis (Figure S6 and Table S2). All signals are predominantly detected in infected regions, indicated by red squares. Scanned images of infected and control samples cross-sections are shown in top row of the figure. Color bar shows intensity of detected ions with yellow and blue colours corresponding to higher and lower intensity, respectively. All images are generated with ± 5mDa error interval.

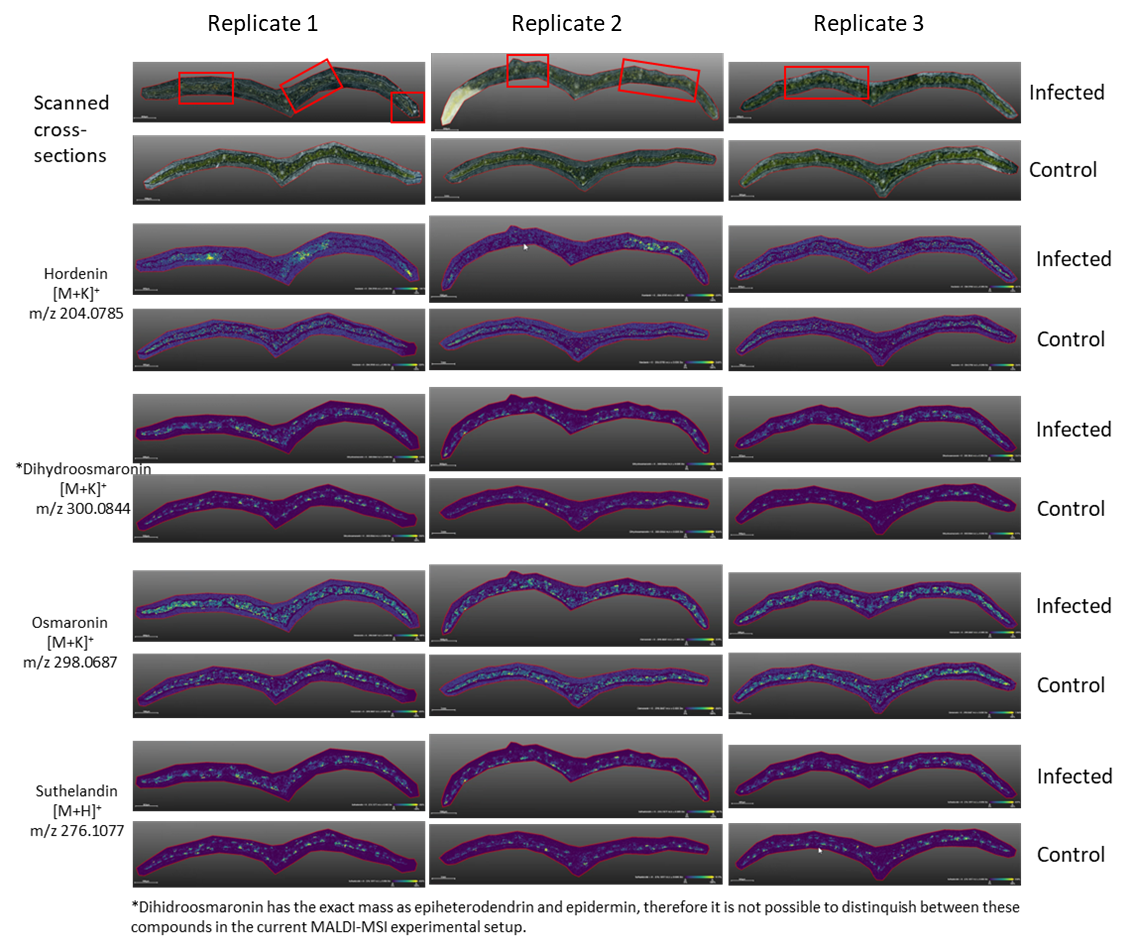

**Figure S9: Localization of phytoanticipins hordenine ([M+K]^+^, *m/z* 204.0785), dihydroosmaronin ([M+K]^+^, *m/z* 300.0844), osmaronin ([M+K]^+^, *m/z* 298.0687) and sutherlandin ([M+H]^+^, *m/z* 276.1077) in infected and control samples of barley leaves using MALDI-MSI.** Compounds are dominantly detected in epidermal reagions of the analyzed barley leaves cross-sections, with exemption of hordenine, dominantly located in the mesophyll of infected regions. Scanned images of infected and control samples cross-sections are shown in top row of the figure. Color bar shows intensity of detected ions with yellow and blue colours corresponding to higher and lower intensity, respectively. All images are generated with ± 5mDa error interval.

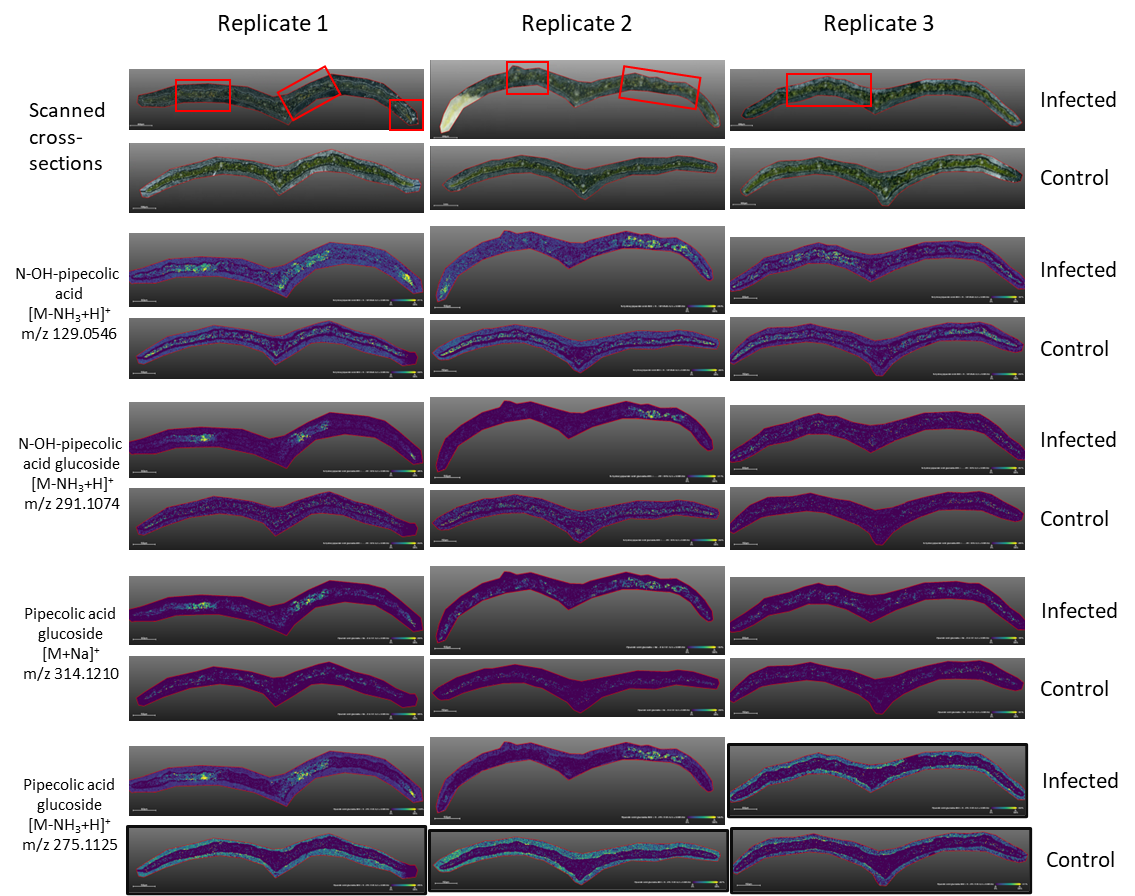

**Figure S10: Localization of *N*-hydroxy pipecolic acid ([M-NH_3_+H]^+^, *m/z* 129.0546), *N*-hydroxy pipecolic acid glucoside ([M-NH_3_+H]^+^, *m/z* 291.1074) and pipecolic acid glucoside ([M+Na]^+^, *m/z* 314.1210; ([M-NH_3_+H]^+^, *m/z* 275.1125)) in cross-sections of infected and control samples of barley leaves using MALDI-MSI with spatial resolution of 20µm.** In the case of images with black border: the lack of signal is most likely caused by the low amount of the compounds in the analyzed replicate; Detection outside of tissue is most likely caused by matrix artefact, considering the uniform distribution pattern. Scanned images of infected and control samples cross-sections are shown in top row of the figure. Color bar shows intensity of detected ions with yellow and blue colours corresponding to higher and lower intensity, respectively. All images are generated with ± 5mDa error interval.

**Table S3: 2OTS orthologues gene copy numbers across barley pangenome** (Jayakodi et al., 2024)

| **Cultivar** | **2OTS gene copies** | **Annuality** | **Origin** | **Row Type** | **Status** | **Subspecies** |
| --- | --- | --- | --- | --- | --- | --- |
| BONUS | 8 | spring | SWE | 2-rowed | cultivar | *vulgare* |
| BOWMAN | 8 | spring | USA | 2-rowed | cultivar | *vulgare* |
| FOMA | 8 | spring | SWE | 2-rowed | cultivar | *vulgare* |
| HID144 | 8 | facultative | IRN | 2-rowed | wild | *spontaneum* |
| HOR 12184 | 8 | winter | CHE | 2-rowed | landrace | *vulgare* |
| HOR 14061 | 8 | spring | TUR | 2-rowed | landrace | *vulgare* |
| HOR 14273 | 8 | spring | SUN |  | landrace | *vulgare* |
| HOR 8148 | 8 | spring | TUR | 2-rowed | landrace | *vulgare* |
| HOCKETT | 7 | spring | USA | 2-rowed | cultivar | *vulgare* |
| HOR 13821 | 7 | spring | TUR | 2-rowed | landrace | *vulgare* |
| RGT PLANET | 7 | spring | DEU | 2-rowed | cultivar | *vulgare* |
| WBDC078 | 7 |  | SYR | 2-rowed | wild | *spontaneum* |
| WBDC184 | 7 |  | LBY | 2-rowed | wild | *spontaneum* |
| BARKE | 6 | spring | DEU | 2-rowed | cultivar | *vulgare* |
| HID249 | 6 | winter | IRN | 2-rowed | wild | *spontaneum* |
| HOR 21256 | 6 | spring |  |  | landrace | *vulgare* |
| WBDC133 | 6 |  | LEB | 2-rowed | wild | *spontaneum* |
| B1K-33-13 | 5 | facultative | ISR | 2-rowed | wild | *spontaneum* |
| GOLDEN PROMISE | 5 | spring | GBR | 2-rowed | cultivar | *vulgare* |
| HID112XXI | 5 | facultative | SYR | 2-rowed | wild | *spontaneum* |
| HID251 | 5 | winter | IRN | 2-rowed | wild | *spontaneum* |
| HID84 | 5 | winter | TUR | 2-rowed | wild | *spontaneum* |
| HOR 2830 | 5 | spring | IRN | 2-rowed | landrace | *vulgare* |
| B1K-17-07 | 4 | facultative | ISR |  | wild | *spontaneum* |
| F2327 | 4 |  | IRN | 2-rowed | wild | *spontaneum* |
| GOLDEN MELON | 4 |  | JPN |  | cultivar | *vulgare* |
| HID357 | 4 | winter | TUR | 2-rowed | wild | *spontaneum* |
| HOR 10892 | 4 | spring | GEO | 2-rowed | landrace | *vulgare* |
| HOR 13594 | 4 | spring | YEM | deficiens | landrace | *vulgare* |
| HOR 14121 | 4 | spring | SYR | 2-rowed | landrace | *vulgare* |
| HOR 1702 | 4 | spring | AFG | 6-rowed | landrace | *vulgare* |
| HOR 18321 | 4 | spring | AFG |  | landrace | *vulgare* |
| HOR 21322 | 4 | winter | SYR | 2-rowed | landrace | *vulgare* |
| HOR 3365 | 4 | winter | RUS | 6-rowed | landrace | *vulgare* |
| HOR 3474 | 4 | winter | RUS | 6-rowed | landrace | *vulgare* |
| HOR 7552 | 4 | spring | PAK | 6-rowed | landrace | *vulgare* |
| HOR 9043 | 4 | spring | ETH | 6-rowed | landrace | *vulgare* |
| MAXIMUS | 4 | spring | AUS | 2-rowed | cultivar | *vulgare* |
| OUN333 | 4 | intermediate | NPL | intermedium | landrace | *vulgare* |
| WBDC103 | 4 |  | JOR | 2-rowed | wild | *spontaneum* |
| WBDC237 | 4 |  | JOR | 2-rowed | wild | *spontaneum* |
| WBDC348 | 4 |  | ISR |  | wild | *spontaneum* |
| AIZU 6 | 3 |  | JPN |  | cultivar | *vulgare* |
| CHIKURIN IBARAKI | 3 |  | JPN | 6-rowed | cultivar | *vulgare* |
| HID055 | 3 | facultative | TUR | 2-rowed | wild | *spontaneum* |
| HID101 | 3 |  | SYR | 2-rowed | wild | *spontaneum* |
| HID380 | 3 | winter | CHN | 6-rowed | wild | *agriocrithon* |
| HOR 10096 | 3 | spring | LBY | 6-rowed | landrace | *vulgare* |
| HOR 1168 | 3 | spring | GRC | 6-rowed | landrace | *vulgare* |
| HOR 12541 | 3 |  |  | 2-rowed | wild | *spontaneum* |
| HOR 13663 | 3 | winter | TUR |  | landrace | *vulgare* |
| HOR 13942 | 3 | spring | ESP | 6-rowed | landrace | *vulgare* |
| HOR 19184 | 3 | spring | IND |  | landrace | *vulgare* |
| HOR 2180 | 3 | spring | CSK | 6-rowed | cultivar | *vulgare* |
| HOR 3081 | 3 | winter | POL | 6-rowed | cultivar | *vulgare* |
| HOR 4224 | 3 | winter | JPN | 6-rowed | cultivar | *vulgare* |
| HOR 495 | 3 | spring | TUR | 6-rowed | landrace | *vulgare* |
| HOR 7172 | 3 | spring | NPL | 6-rowed | landrace | *vulgare* |
| HOR 9972 | 3 | spring | PAK | 6-rowed | landrace | *vulgare* |
| IGRI | 3 | winter | DEU | 2-rowed | cultivar | *vulgare* |
| MOREX | 3 | spring | USA | 6-rowed | cultivar | *vulgare* |
| WBDC199 | 3 |  | SYR | 2-rowed | wild | *spontaneum* |
| WBDC207 | 3 |  | UZB | 2-rowed | wild | *spontaneum* |
| ZDM01467 | 3 | spring | CHN | 6-rowed | landrace | *vulgare* |
| ZDM02064 | 3 | spring | CHN | 6-rowed | landrace | *vulgare* |
| 10TJ18 | 2 |  | TJK |  | landrace | *vulgare* |
| AKASHINRIKI | 2 | winter | JPN | 6-rowed | cultivar | *vulgare* |
| B1K-04-12 | 2 | facultative | ISR | 2-rowed | wild | *spontaneum* |
| HOR 10350 | 2 | spring | ETH | 6-rowed | landrace | *vulgare* |
| HOR 21595 | 2 | spring | SYR |  | landrace | *vulgare* |
| HOR 2779 | 2 | spring | IRN | 2-rowed | landrace | *vulgare* |
| HOR 6220 | 2 | spring | ETH | labile | landrace | *vulgare* |
| HOR 7385 | 2 | spring | CZE | 2-rowed | landrace | *vulgare* |
| HOR 8117 | 2 | spring | TUR | 2-rowed | landrace | *vulgare* |
| WBDC349 | 2 |  | ISR |  | wild | *spontaneum* |

| **Table S4:** **Quantification of hydroxylated tryptamine isomers across different barley cultivar and pathogen combinations.** Quantification of serotonin, hydroxylated tryptamine derivative [2], and 2OT in barley leaves from three different cultivars: Pallas P-02, Golden Promise, and RGT Planet infected with powdery mildew (*B. hordei*), *P. syringae*, and *P. teres*. Values are the mean of n = 3 biological replications ± S.E. Abbreviations: 2OT, 2-oxo-tryptamine; FW, fresh weight; S.E., standard error of the mean. | | | | | |
| --- | --- | --- | --- | --- | --- |
| Pathogen | Cultivar | Time point | Mean serotonin [nmol/g FW]  ± S.E. | Mean hydroxylated tryptamine derivative [2]  [nmol/g FW] ± S.E. | Mean 2OT  [nmol/g FW]  ± S.E. |
| Powdery mildew  (*B. hordei*) | Pallas  P-02 | 0 | 0.32 ± 0.02 | 0.31 ± 0.02 | 0.72 ± 0.72 |
|  |  | 2 | 0.30 ± 0.001 | 0.39 ± 0.01 | 7.48 ± 0.76 |
|  |  | 7 | 4.73 ± 2.17 | 1.55 ± 0.51 | 52.54 ± 14.45 |
| *P. syringae* | Golden Promise | 4 | 15.52 ± 2.53 | 23.43 ± 4.79 | 207.34 ± 78.47 |
|  |  | 7 | 19.87 ± 4.15 | 18.47 ± 6.24 | 210.58 ± 78.21 |
|  | RGT Planet | 4 | 11.17 ± 1.13 | 18.66 ± 0.25 | 224.58 ± 2.42 |
|  |  | 7 | 21.97 ± 4.70 | 17.46 ± 2.31 | 178.88 ± 30.21 |
| *P. teres* | Golden Promise | 4 | 8.01 ± 4.45 | 1.98 ± 0.70 | 3.86 ± 3.86 |
|  |  | 7 | 1.93 ± 0.81 | 0.97 ± 0.16 | 0 ± 0 |
|  |  | 12 | 1.61 ± 0.56 | 0.82 ± 0.1 | 0 ± 0 |
|  | RGT Planet | 4***** | 19.39 ± 1.16 | 5.76 ± 0.89 | 32.66 ± 6.12 |
|  |  | 7****** | 66.26 ± 29.96 | 9.29 ± 3.50 | 31.40 ± 25.93 |
|  |  | 12 | 43.03 ± 3.90 | 38.20 ± 16.31 | 60.57 ± 4.20 |
| ***** n = 4, ****** n = 2. | |  |  |  |  |

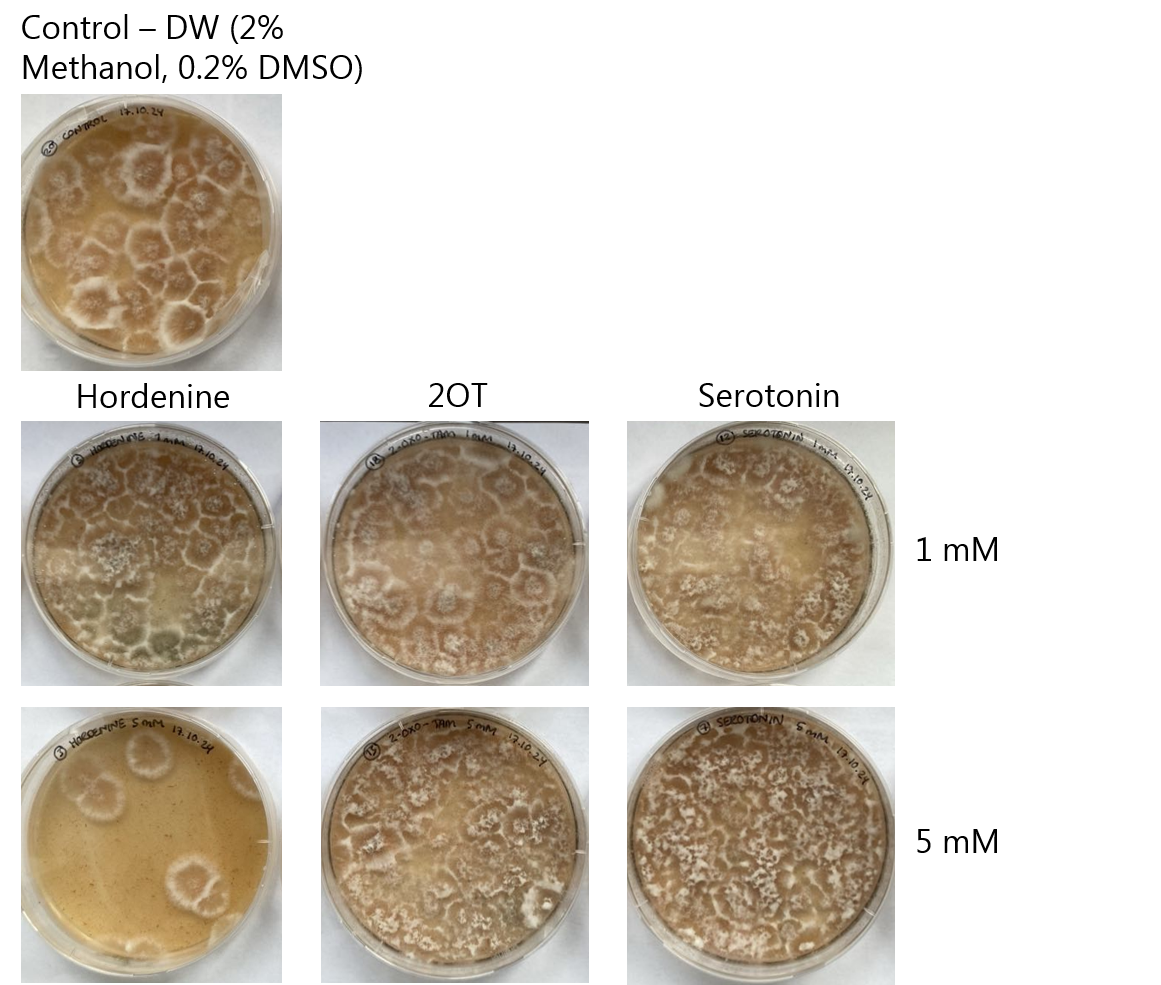
**Figure S11:** **Evaluation of *P. teres* fungal growth after five days incubation.** Spores of *P. teres* were incubated with 1 or 5 mM hordenine, 2OT, or serotonin for 24 hours, plated and incubated for five days at RT. Demineralized water (2% methanol, 0.2% DMSO) was used as control. 2OT and serotonin showed no inhibition compared to hordenine, which at 5 mM demonstrated clear inhibitory effect on fungal growth. Abbreviations: DW, demineralized water; DMSO, dimethyl sulfoxide; 2OT, 2-oxo-tryptamine; RT, room temperature.

**Figure S12: Visual representation of annotated chemical space in barley leaves after infection by *P. teres*. (A)** Global feature-based molecular network (FBMN) visualizing the detected chemical space of barley *cv.* RGT Planet. The 38 largest molecular families (i.e. ≥ 3 nodes) in the network are numbered. In this study, 16.5% of 922 features were annotated (highlighted in green). List of annotated compounds are found in Table S1. **(B)** Selected molecular families from network, which are induced in infected samples, but remain completely unannotated. At the top, pie-charts indicate relative abundance of individual features in control (blue) and infected (orange) samples across entire spectral dataset. Below, nodes are coloured according to compound class (NPC Pathway) predictions from SIRIUS. Abbreviations: NPC, NPClassifier taxonomy (Kim et al., 2021).

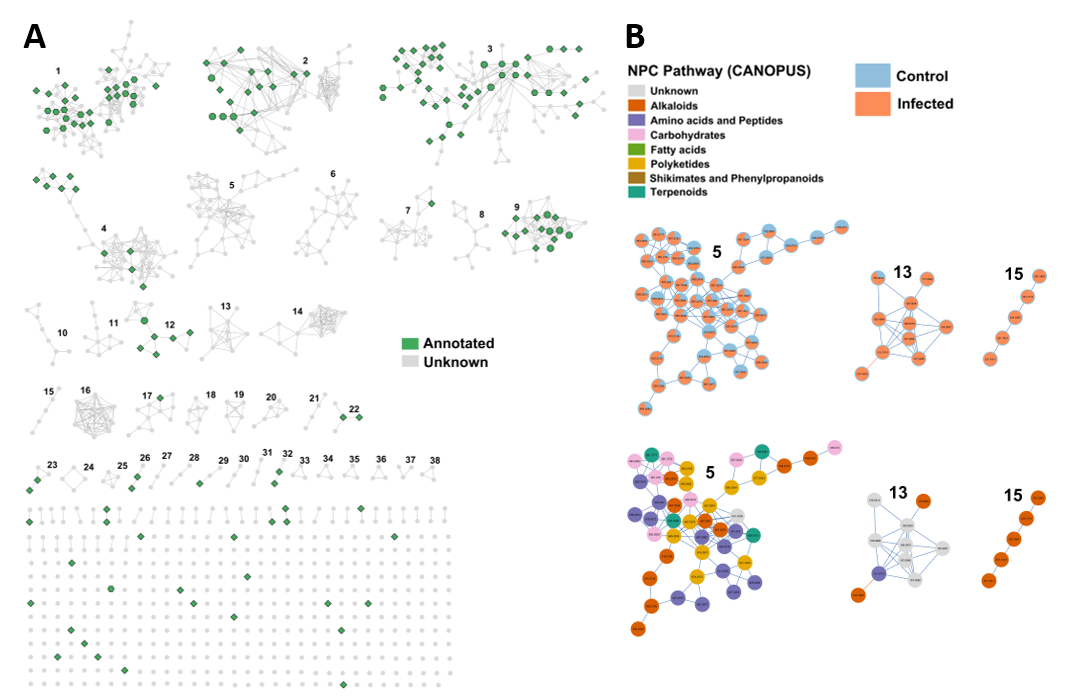

| **Table S5: Parameters for data processing of spectral data in MZmine4, GNPS2 (FBMN), and SIRIUS.** | | |
| --- | --- | --- |
| **MZmine4 (v. 4.0.3)** | | |
| **Step** | **Parameter** | **Value** |
| Mass detection | Mass detector | Centroid |
|  | Polarity | + |
|  | Scan types (IMS) | All scan types |
|  | MS level | 1 |
|  | Noise level | 2.0E2 |
| Mass detection | Mass detector | Centroid |
|  | Polarity | + |
|  | Scan types (IMS) | All scan types |
|  | MS level | ≥ 2 |
|  | Noise level | 1.0E2 |
| ADAP Chromatogram Builder | MS level | 1 |
|  | Retention time | 0.50-15.00 min. |
|  | Polarity | + |
|  | Spectrum type | Centroid |
|  | Min consecutive scans | 5 |
|  | Min intensity for consecutive scans | 6.0E2 |
|  | Min absolute height | 2.0E3 |
|  | m/z tolerance | 0.005 m/z or 20 ppm |
| Smoothing | Smoothing algorithm | Savitzky-Golay |
| Chromatogram Resolving | Algorithm | Local minimum feature resolver |
|  | Chromatographic threshold | 90% |
|  | MS/MS scan pairing | YES |
|  | Min in RT range | 0.07 min |
|  | Min relative height | 0.0% |
|  | Min absolute height | 2.0E3 |
|  | Min ratio of peak top/edge | 2.0 |
|  | Peak duration range | 0.00 – 1.51 min |
|  | Minimum scans (data points) | 5 |
|  | MS2 pairing: m/z range | 0.01 m/z or 20.0 ppm |
|  | MS2 pairing: RT range | Feature edges |
| 13C isotope filter | m/z tolerance | 0.01 m/z or 10 ppm |
|  | RT tolerance | 0.1 min |
|  | Monotonic shape | YES |
|  | Max charge | 2 |
|  | Representative isotope | Most intense |
|  | Never remove feature with MS2 | YES |
| Isotopic peaks finder | Chemical elements | H, C, N, O, S |
|  | m/z tolerance (feature-to-scan) | 0.001 m/z or 5 ppm |
|  | Max charge of isotope m/z | 2 |
|  | Search in scans | Single most intense |
| Join aligner | m/z tolerance | 0.006 m/z or 10 ppm |
|  | Weight for m/z | 3 |
|  | RT tolerance | 0.1 min |
|  | Weight for RT | 1 |
|  | Mobility weight | 1.000 |
| Feature list rows filter | Min aligned features (samples) | 2 samples or 10% |
|  | Validate 13C isotope pattern | YES |
|  |  | m/z tolerance: 0.0015 m/z or 3 ppm |
|  |  | Max charge: 2 |
|  |  | Estimate minimum carbon: YES |
|  |  | Remove if 13C: YES |
|  |  | Exclude isotopes: O |
|  | Keep or remove rows | Keep rows that match all criteria |
|  | Never remove feature with MS2 | YES |
| Gap filling | Algorithm | Peak finder (multithreaded) |
|  | Intensity tolerance | 20% |
|  | m/z tolerance (sample-to-sample) | 0.005 m/z or 20 ppm |
|  | Retention time tolerance | 0.1 min |
|  | Minimum scans (data points) | 3 |
| Duplicate peak filter | Filter mode | New average |
|  | m/z tolerance | 0.0008 m/z or 1.5 ppm |
|  | RT tolerance | 0.04 min |
| Correlation grouping  (metaCorrelate) | RT tolerance | 0.05 min |
|  | Min feature height | 0 |
|  | Intensity threshold for correlation | 5.0E2 |
|  | Feature shape correlation | YES |
|  | Feature height correlation | YES |
| Ion identity networking | m/z tolerance | 0.0015 m/z or 3 ppm |
|  | Check | All features |
|  | Min height | 0 |
|  | Ion identity library (adducts  and modifications) | [M+H]+ (1.0073)  [M+Na]+ (22.9892)  [M+K]+ (38.9632)  [M+NH4]+ (18.0338)  [M+2H]2+ (2.0146)  [M-H+2Na]+ (44.9712)  [M-H2O] (-18.0106)  [M-2H2O] (-36.0211) |
|  | Annotation refinement | YES |
| Spectral library search | MS level | ≥ 2 (merged) |
|  | Precursor m/z tolerance | 0.01 m/z or 20 ppm |
|  | Spectral m/z tolerance | 0.01 m/z or 20 ppm |
|  | Remove precursor | YES |
|  | Min matched signals | 4 |
|  | Similarity | Weighted cosine |
|  | Weights | SQRT (mz^0*\|^0.5) |
|  | Min cosine similarity | 0.7 |
|  | Handle unmatched signals | Keep all and match to zero |
| MS/MS spectral networking | m/z tolerance (MS2) | 0.005 m/z or 20 ppm |
|  | Only best MS2 scan | YES |
|  | Max precursor m/z delta | 500 |
|  | Min matched signals | 4 |
|  | Min cosine similarity | 0.7 |
| Export spectral networks to graphml (FBMN/IIMN) | NA | NA |
| Export molecular networking files (e.g., GNPS, FBMN, IIMN, MetGem) | Filter rows | MS2 or ion identity |
|  | Feature intensity | Area |
|  | CSV export | Simple |
|  | Merge MS/MS | YES |
|  | Select spectra | Across samples |
|  | m/z merge mode | Weighted average (remove outliers) |
|  | Intensity merge mode | Sum intensity |
|  | Exp. mass deviation | 0.01 m/z or 15 ppm |
|  | Cosine threshold | 70% |
|  | Peak count threshold | 20% |
|  | Isolation window offset (m/z) | 0 |
|  | Isolation window width (m/z) | 3 |
| Export for SIRIUS | m/z tolerance | 0.0020 m/z or 5 ppm |
|  | Merch MS/MS | YES |
| Export all annotations to CSV file | Top N per method | 10 |
| **GNPS2 (Release 2024.09.17): Workflow for feature-based molecular networking (v. 2024.10.09)** | | |
| **Step** | **Parameter** | **Value** |
| General parameters | Precursor Ion Tolerance | 0.02 Da |
|  | Fragment Ion Tolerance | 0.02 Da |
| Advanced Filtering parameters | Min Peak Intensity | 0.0 |
|  | Window Filter | Yes |
|  | Precursor Window Filter | Yes |
| Networking Parameters | min_cosine | 0.7 |
|  | min_matched_peaks | 6 |
|  | networking_max_shift | 1999 |
| Network Topology Parameters | Classic Param – Top K | 10 |
|  | Classic Param – Max Component Size | 100 |
| Library Search Paramenters | Library Minimum Cosine | 0.7 |
|  | Library Minimum Matched Peaks | 6 |
|  | Analog Search | No/Yes |
|  | Analog Search Max Shift | 1999 |
|  | Top-K | 1 |
| Normalization Parameters | Quantification Normalization | None |
| **SIRIUS (v. 5.8.6)*** | | |
| **Step** | **Settings** | |
| SIRIUS | Q-TOF is selected as instrument used for analysis, i.e. MS2 mass accuracy is set to 10 ppm. Remaining settings are left as default. | |
| CSI:FingerID | ‘Search DBs’ is enabled (Bio Database and Natural Products). Remaining settings are left as default. Fallback adducts: [M + H]+, [M + K]+, [M + Na]+, [M + H3N + H]+, [M + CH4O + H]+, [M + C2H3N + H]+. | |
| ZODIAC | Settings are left as default. | |
| CANOPUS | Parameter-free. | |
| *Compounds with a mass above 850 Da were excluded from analysis due to computational limits | | |

| **Table S6: List of authentic standards used as LC-MS/MS in-house spectral library.** Structural information on all reference compounds used in this study. | | | |
| --- | --- | --- | --- |
| **Compound** | **Structure** | **Formula** | **Monoisotopic mass** |
| 2-oxo-tryptamine (2OT) | 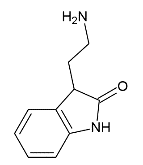 | C10H12N2O | 176.09495 |
| Tryptamine | 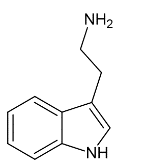 | C10H12N2 | 160.10004 |
| Epiheterodendrin | 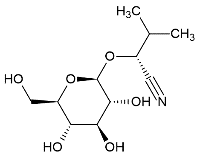 | C11H19NO6 | 261.12122 |
| Epidermin | 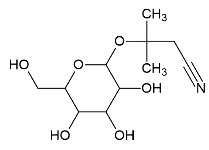 | C11H19NO6 | 261.12122 |
| Osmaronin | 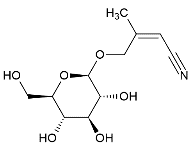 | C11H17NO6 | 259.10557 |
| Dihydroosmaronin | 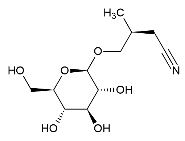 | C11H19NO6 | 261.12122 |
| Sutherlandin | 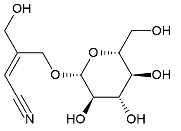 | C11H17NO7 | 275.10049 |
| 3-(2-aminoethyl)-3-hydroxy-indolin-2-one (AEHI) | 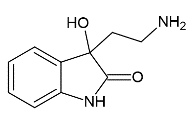 | C10H12N2O2 | 192.08987 |
| Serotonin | 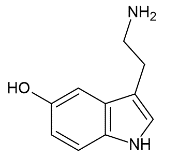 | C10H12N2O | 176.09495 |
| Tryptophan | 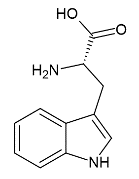 | C11H12N2O2 | 204.08987 |
| Hordenine | 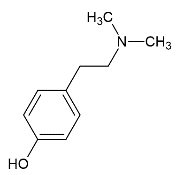 | C10H15NO | 165.11536 |
